# Polystyrene microplastic uptake drives Inflammatory, Epitranscriptomic, and Metabolic Reprogramming in Human Aortic Endothelial cells

**DOI:** 10.64898/2026.07.09.737624

**Authors:** Ajmal Khan, Grant Koher, Tariq Khan, Kyla Grant, Sarah Hudson, Gordon Zheng, Ho Young Lee, Warren S. Vidar, Frank Morales-Shnaider, Jinlan Chen, Kwame A. Darfour-Oduro, Ramji Bhandari, Xuewei Zhu, Kerui Wu, Norman Chiu, Zhenquan Jia

## Abstract

Microplastics (MPLs) are pervasive environmental pollutants increasingly linked to adverse human health outcomes, including atherosclerosis. However, the underlying mechanisms remain poorly understood. Human aortic endothelial cells (HAECs), which line the inner surface of blood vessels, play a critical role in maintaining vascular homeostasis and in the development of atherosclerosis. This study demonstrates that polystyrene microplastics enter HAECs through clathrin-mediated endocytosis and macropinocytosis and subsequently co-localize with mitochondria and lysosomes. Exposure to MPLs induced coordinated transcriptional, epitranscriptomic, and metabolomic reprogramming in HAECs. Transcriptomic analysis revealed disruption of mitochondrial genes and activation of inflammatory pathways with the response of the NF-κB pathway being particularly prominent. Mass spectrometry analysis of RNA modification further identified significant remodeling of the epitranscriptomic landscape, highlighted by increased 1-methyladenosine (m1A) modification and reciprocal regulation of its associated enzymes (*TRMT61A* upregulation and *ALKBH3* suppression), along with alterations in other RNA modifications such as m3C, pseudouridine (Ψ), m5C, and m7G. Comparative analysis of transcriptomic profiles from human atherosclerotic plaques revealed shared dysregulated pathways in vascular regulation and cellular signaling. Metabolomic profiling further showed extensive remodeling of lipid metabolic networks associated with oxidative stress and inflammation. Together, these findings suggest that MPLs exposure may disrupt endothelial function and pose a potential risk to human cardiovascular health.

**Graphical Abstract:** 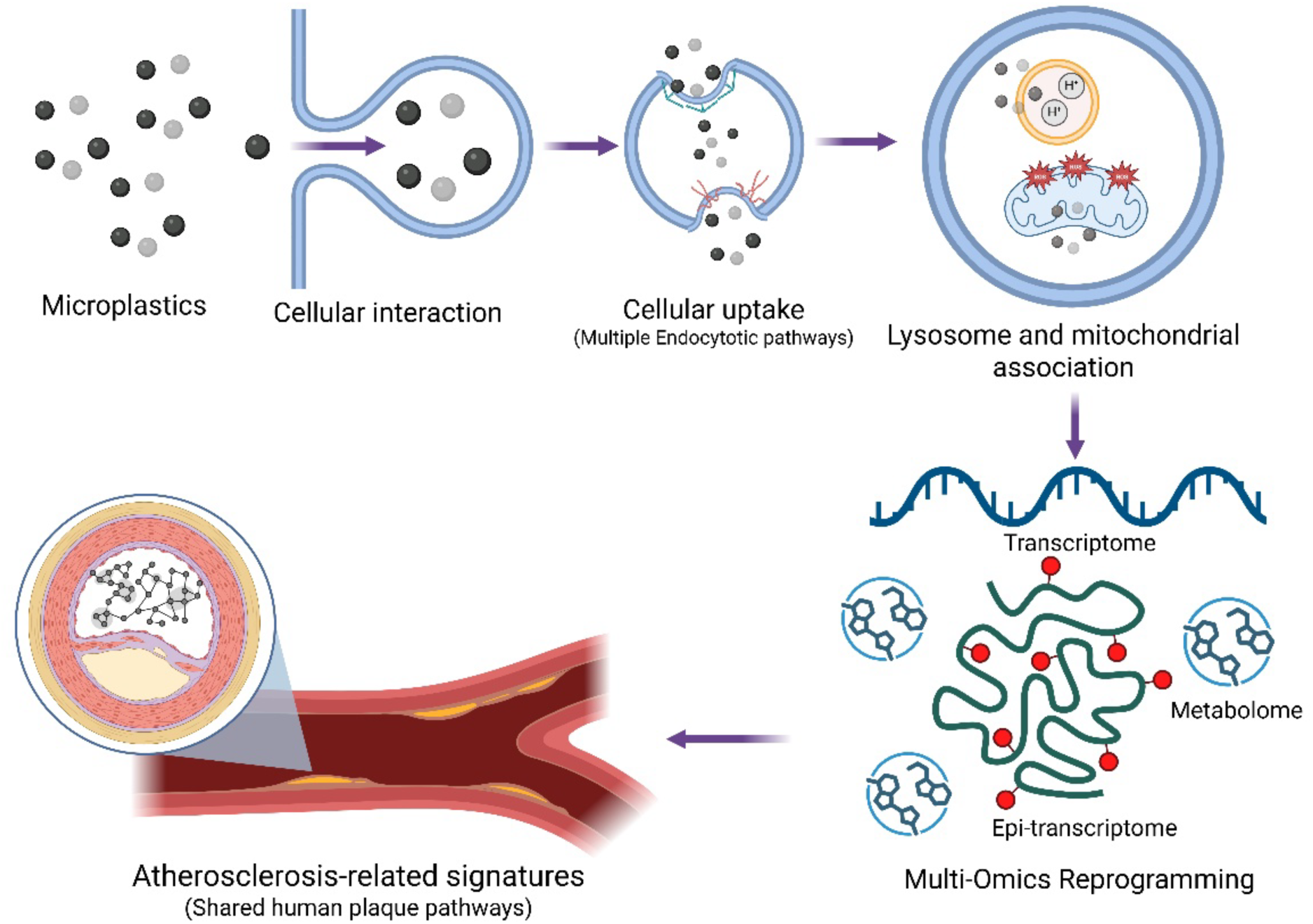

## 1. Introduction

Microplastics have been recognized as a novel environmental pollutant and are considered an emerging threat to ecosystems and human health. These plastic particles, commonly found in the environment, have been detected in human food, plants, water, and air. They are the result of environmental degradation, chemical breakdown, or mechanical wear and tear ^1^. Microplastics can enter the body through ingestion, inhalation, and dermal contact ^2^, and have been detected in human blood ^3^, stool ^4^, placenta ^5^, lungs ^6^, sputum ^7^, saliva, hair, facial skin, hand skin ^8^, as well as in the carotid artery plaque of human patients ^9^ and the pericardial sac of zebrafish ^10^. Various types of microplastics (polystyrene, polyurethane, polyethylene, polyoxymethylene, polyamide, etc.) exist in the environment ^11^, among which polystyrene (PS) is one of the abundant and commonly detected plastic particles ^12–17^. The toxic effects of environmental exposure to microplastics have been reported in experimental animal models, cell lines, and wild animal species, including aquatic and terrestrial species^18–20^. It is envisioned that if left unchecked, microplastics will continue to grow as a significant problem, leading to major economic and health issues. This is evidenced by the fact that microplastics are cytotoxic to a variety of human cell lines, such as GES-1 cells ^18^, Caco-2 cells ^19^, and HEK293 ^21^, and have been associated with ovarian fibrosis ^22^, pulmonary fibrosis, and tissue injury in the kidney, liver, intestine, brain, and testis ^19^.

Cardiovascular diseases (CVDs) are the leading cause of death worldwide ^23^. Each year, approximately 20.5 million people die from CVDs, accounting for approximately 33% of all global deaths ^23^. Atherosclerosis is a major chronic inflammatory form of CVD, characterized by the thickening of blood vessels caused by the build-up of plaque (deposits of fatty substances, cholesterol, cellular waste products, calcium, and fibrin) in the inner lining of blood vessels ^24, 25^. Among the various risk factors of atherosclerosis, environmental microplastics and emerging environmental contaminants have recently been recognized as potentially important contributors to human atherosclerotic cardiovascular disease. Recently, a prospective cohort study reported that human carotid artery atheroma contained microplastics and nanoplastics, predominantly polyethylene (mean 21.7 ± 24.5 µg per mg of plaque), and that their presence was associated with a 4.5-fold higher risk of developing various cardiovascular events (myocardial infarction, stroke, or death), providing the first clinical evidence that suggests microplastics and nanoplastics are risk factor for human cardiovascular events ^9^. Notably, studies found that exposure to microplastics caused cardiac fibrosis and apoptosis of the myocardium ^26^. The endothelium, which lines the interior of the heart and blood vessels, is a prominent target of environmental contaminants ^27, 28^. Microplastics were shown to increase endothelial cell peroxynitrite levels, leading to oxidative injury ^29^.

Although microplastics have been detected in human organs, including the cardiovascular system, their uptake pathways and transcriptomic and metabolomic effects in human endothelial cells remain unclear. This study explored the uptake and internalization mechanisms of polystyrene microplastics (MPLs) in human aortic endothelial cells (HAECs) and further analyzed whether MPLs-treated HAECs recapitulate the characteristics of human atherosclerosis at the transcriptomic, epitranscriptomic, and metabolomic levels. The results showed that HAECs take up MPLs through clathrin-mediated endocytosis and macropinocytosis. MPLs induced an inflammatory response, specifically activating the TNF-α and NF-κB signaling pathways. Additionally, MPLs treatment altered HAECs metabolic profile, including metabolites implicated in pathways relevant to cardiovascular and inflammatory disease. Simultaneously, MPLs exposure led to abnormalities in various RNA modifications, suggesting altered expression of related genes. Comparison of the transcriptomic profiles of MPLs-treated HAECs and human atherosclerotic plaques revealed common dysregulation across several pathways, particularly those involved in vascular physiological regulation and cell signaling. Our study reveals that HAECs can internalize MPLs, leading to dysregulation of transcriptomic, epitranscriptomic, and metabolic pathways, suggesting that MPLs exposure may have potential adverse effects on human health.

## 2. Methods and Materials

### 2.1. Cell culture

Primary human aortic endothelial cells (HAECs; ATCC) were cultured in endothelial MCDB-131 medium supplemented with 10% fetal bovine serum (FBS), 10 mM L-glutamine, 1% penicillin/streptomycin, 1 µg/mL hydrocortisone, and 10 ng/mL epidermal growth factor. Cells were maintained in Cellstar® Filtered Cap 75 cm^2^ cell–culture treated screw cap flasks in incubators set to 37 °C and 5% CO_2_. Subculturing was performed when cells reached 85–95% confluence. An aliquot was then transferred to Corning® 12 Well Cell Culture Plates and grown to 80% confluence before treatment. For the uptake and microscopy experiments, cells were treated with 80 nm sized Fluorescent labeled Sphero™ Polystyrene particles (MPLs; Sphero™ Cat No.: CFP-00552-2, 10mg/mL) and for other experiments, cells were treated with 80 nm sized non-Fluorescent labeled Sphero™ Polystyrene particles (MPLs; Sphero™ Cat No.: PP-008-10, 50 mg/mL). Both polystyrene particles were supplied as an aqueous suspension containing 0.02% (w/v) sodium azide. All microplastic treatment concentrations were prepared from corresponding stocks and added at the same volume, each treatment contained the same final concentration of sodium azide (0.00005%); therefore, a matched sodium azide vehicle control was used for all treatment groups.

### 2.2. Uptake, cell viability and routes of MPLs internalization

Cells were cultured in 12-well plates with endothelial cell culture medium to 85% confluence. The cells were then treated with 0 µg/mL (Ctrl group) and 10, 20, and 120 µg/mL polystyrene microplastics (MPLs group) for 24 h. During treatment, plates were incubated at 37 °C and 5% CO_2_. After treatment, cells were rinsed with 1× Phosphate Buffer Solution (PBS) twice and collected using cell scraping in 600 µL 1× PBS. The collected cells were aliquoted into a black 96-well Costar plate at 200 µL per well. The fluorescence was read using a Bio-Tek® Synergy 2™ plate reader at 360/40 excitation and 460/40 emission. Samples were then analyzed by flow cytometry using a Guava® easyCyte™ flow cytometer (Millipore™, GuavaSoft 3.1.1) using the InCyte assay. The percentage of GFP-positive cells was quantified as the percentage of events within the R1 region set on the Green-B (GRN-B-HLog) fluorescence histogram. Cell viability was determined at 0, 10, and 120 µg/mL using slight modified version of MTT reduction assay ^30, 31^.

To determine the routes of MPLs internalization, cells were washed twice with 1× PBS and then incubated with varying concentrations of channel blockers and endocytic inhibitors **(Table S1)** for 30 min ^32^. After 30 minutes, cells were washed twice with 1× PBS and then incubated with MPLs (10 µg/mL) for 24 h. Following completion of the incubation period, the medium and inhibitors were removed, and the cells were rinsed twice with 1× PBS. Cell collection was performed using a cell scraping method in 600 µL of 1× PBS. The collected cells were aliquoted into a black 96-well Costar plate at 200 µL per well. The plate fluorescence was read using a Bio-Tek® Synergy 2™ plate reader at 360/40 excitation and 460/40 emission. Samples were also analyzed by flow cytometry using a Guava® easyCyte™ flow cytometer (Millipore™) set to Green-B fluorescence, and the percentage of GFP-positive cells was quantified as described above.

### 2.3. Fluorescence Microscopy

HAECs were treated with MPLs (10 µg/mL) of 80 nm MPLs for 24 h. The Keyence BZ-X710 All-in-One Fluorescence Microscope was used to visualize the localization of MPLs in HAECs. Representative Phase-contrast, GFP (green), and overlay images indicated that MPLs were taken up by HAECs. Imaging was performed with n=3 wells/condition and 9 fields of view were acquired/experiment.

### 2.4. MPLs–Mitochondria Co-localization

HAECs were pre-incubated with 0.5 μM of MitoTracker™ Red CMXRos for 30 min and then treated with MPLs (Size: 80 nm, Concentration: 10 µg/mL) for 24 h. The Keyence BZ-X710 All-in-One Fluorescence Microscope was used to visualize the interaction of MPLs with mitochondria in HAECs. Representative phase-contrast (PC), GFP (green), Cy5 (red), and overlay (PC/GFP, Cy5/GFP) images showed that MPLs interacted with endothelial mitochondria. Co-localization imaging was performed with n=3 wells/condition and with 9 fields of view/condition.

### 2.5. MPLs-Lysosome Co-localization

HAECs treated with MPLs were incubated with LysoTracker Deep Red (Thermo Fisher Scientific, Cat. No. L12492) at a concentration of 100 nM for the last 2 h of the 24 h MPLs exposure period. Following the incubation period, the old media (containing MPLs and LysoTracker Deep Red dye) were removed, the cells were washed with PBS three times and then supplemented with fresh media. Nikon confocal microscope (AXR inverted dual point-scanning confocal/widefield microscope equipped with the AXR 2k Resonant Confocal scanner) was used to acquire images with 40x air objective (CFI60 Plan Apo Lambda D; NA 0.95; correction collar 0.17–0.25 mm; FOV 25 mm; DIC; spring-loaded). Cells were maintained in a live-cell environment (37°C and 5% CO₂) throughout image acquisition. LysoTracker Deep Red fluorescence was imaged with excitation at ∼647 nm and emission at ∼668 nm. Fiji ImageJ was used for image processing and analysis (merging images).

### 2.6. Luciferase Assay

NF-κB Luciferase Reporter HEK293 Stable Cell Line (SL-0012-NP) was purchased from Signosis, Inc. (Santa Clara, California). The cells were cultured in DMEM medium with 10% FBS and 1% penicillin-streptomycin. NF-κB reporter cells (1 × 10^4^) plated in a 96-well white-wall plate were treated with MPLs (120 µg/mL). Following incubation, the media was removed, cells were washed with 1× PBS and then lysed for 15 minutes with 20 µl of 1x lysis buffer. Following this, 100 µL of luciferase substrate was added, mixed, and read on a luminometer.

### 2.7. RNA Sequencing

HAECs were treated with MPLs (size: 80 nm, concentration: 120 µg/mL) in 12-well plates at 85% confluency in complete MCDB-131 medium and incubated for 24 h at 37 °C and 5% CO_2_. RNA was extracted using the Quick-RNA Microprep Kit (#R1051, Zymo Research, CA, USA) according to the manufacturer’s protocol ^33^. RNA quality was determined using Nanodrop 2000 and Qubit (Thermo Fisher, Waltham, MA). RNA from three wells was pooled to form a single biological replicate, and RNA libraries were prepared using the NEBNext® Ultra II RNA ^TM^ II RNA Library Prep Kit for Illumina®. The prepared libraries were visualized and tested for quality with Gel Electrophoresis. The prepared libraries were sequenced using Illumina HiSeq X (Novogene Corporation, CA, USA) using a 150 bp paired-end sequencing strategy (short reads) and produced 50–95 million reads per sample.

The reads were first preprocessed with fastp (v 0.23.2) ^34^, an ultra-fast all-in-one FASTQ preprocessor that performs adapter trimming, quality filtering, and per-read quality pruning with a single scan of the FASTQ data. The processed reads were then mapped to the human genome (hg38 analysis set) using the STAR (v2.7.7a) RNA-seq aligner ^35^. Gencode reference annotation for human (v42) ^36^ was used for annotation. Finally, DESeq2 (v1.34.0) ^37^ was used for differential expression analysis. Genes with log_2_ fold change ≥ 1 or ≤ -1 and a q value ≤ 0.05 were considered differentially expressed. For post-analysis of our data, we used online platforms, including: 1) https://bioinformatics.sdstate.edu/idep20/ ^38^, 2) https://bioinforx.com/web/product_bxtoolbox.php ^39^, 3) https://bioinformatics.sdstate.edu/go/ ^40^, 4) https://xomicsshiny.bxgenomics.com/ ^41^. DEGs were manually curated into upregulated pro-atherosclerotic and downregulated anti-atherosclerotic genes. To determine the effects of MPLs on mitochondria, mitochondrial DEGs (padj < 0.05) were identified **(Fig. 2d)**.

### 2.8. Integrative Transcriptomics Approach to demonstrate Molecular determinants of MPLs uptake

To identify molecular pathways involved in MPLs internalization, pathway and functional enrichment analyses were performed on 43 differentially expressed genes (DEGs) involved in cellular uptake and vesicle trafficking. Gene Ontology (GO) enrichment analysis was performed for Biological Processes (BP) and Cellular Component (CC) using the (https://bioinformatics.sdstate.edu/idep20/) platform, and any terms with padj value < 0.05 were considered significantly enriched. BP terms related to MPLs internalization, such as endocytosis and vesicle-mediated transport, and CC terms related to localization of MPLs, such as caveolae, early endosome, and vesicle membrane, were prioritized. For further resolution of MPLs uptake pathways, we performed Kyoto Encyclopedia of Genes and Genomes (KEGG) pathway enrichment analysis on significant DEGs using (https://bioinformatics.sdstate.edu/idep20/). The KEGG endocytosis pathway map was generated to visualize gene-level enrichment in endocytosis. Gene involved in various internalization mechanisms, such as clathrin-mediated and clathrin-independent endocytosis, were identified by overlaying DEGs onto the KEGG endocytosis pathway (hsa04144). To visualize the expression patterns of selected DEGs involved in MPLs internalization, hierarchical clustering was performed using Euclidean distance, and a heatmap was generated. The functional annotations of the heatmap visualized DEGs involved in the internalization of MPLs were cross-checked to confirm pathway-level association.

### 2.9. Comparative transcriptomic analysis between MPLs-treated HAECs and Human Atherosclerotic plaques

To compare gene expression profiles between atherosclerotic plaques and MPLs-treated HAECs, publicly available bulk RNA sequencing data (GSE120521) of stable and unstable human atherosclerotic plaques were retrieved from NCBI Gene Expression Omnibus (GEO). DEGs from publicly available datasets and MPLs-treated HAECs were compared using (https://bioinforx.com/web/product_bxtoolbox.php), yielding common and uniquely expressed DEGs. To assess regulatory concordance between DEGs, common DEGs were further classified into mutually upregulated, mutually downregulated, and discordantly regulated DEGs based on fold change values. To determine the role of common DEGs, pathway enrichment analysis was performed on the common DEGs (n = 457) by subjecting common DEGs to KEGG pathway enrichment analysis (https://bioinformatics.sdstate.edu/go/). Fold enrichment and - log_10_(FDR) values were used to quantify pathway significance.

### 2.10. RNA modification

To determine the epitranscriptomics changes induced by MPLs, we quantified multiple modified ribonucleosides in total RNA isolated from HAECs. To properly assess epitranscriptomics roles of MPLs in HAECs, cells were treated at 80% confluence with MPLs at concentration of 0 and 120 µg/mL. Following incubation, the Quick-RNA™ MicroPrep kit was used to extract RNA. The epitranscriptome analysis was performed using UPLC–MS. The data analysis was performed using ionization efficiency normalization based on the SqEP method ^42^. Each RNA sample was enzymatically digested in a 25 μL reaction at 37 °C for 3 h. The reaction mixture included 5 μg of RNA, 0.05 units of Phosphodiesterase I, 0.5 units of Alkaline Phosphatase, 5 units of Benzonase, 50 mM Tris-HCl (pH 8.0), 1 mM MgCl_2_, and 0.1 mg/mL BSA.

Following digestion, UPLC-MS Analysis: Chromatographic separation of ribonucleoside standards from digested RNA samples was performed on an Acquity ultra-performance liquid chromatography (UPLC) system (Waters Corporation, Milford, MA) with a binary pump and an autosampler maintained at 4 °C. A Water’s Acquity UPLC HSS T3 column (2.1 x 50 mm, 1.8 µm particle size) and a HSS T3 VanGuard precolumn (2.1mm x 5 mm, 1.8 µm particle size) with an Acquity inline filter kit held at 30 °C were used. Unless otherwise stated, 10 µL of 50 ng/µL of digested RNA sample was injected onto the column, and the elution was carried out with a two-solvent system prepared with Optima grade solvents. Mobile Phase A consisted of H^2^O with 0.01% formic acid, and Mobile Phase B consisted of 50% CH_3_CN with 0.01% formic acid held at a flow rate of 0.4 mL/min. The binary elution gradient was carried out as follows: An initial isocratic composition of 100:0 (A:B) from 0.0 – 0.5 min., increasing to 70:30 from 0.5 – 9.0 min. at a curve of 7, linearly changing to 50:50 from 9.0 - 10.0 min., followed by another linear increase to 0:100 from 10.0 – 15.0 min., a final isocratic hold at 0:100 from 15.0 – 16.0 min. marks the transition to column wash and re-equilibration at the starting conditions for 8 minutes. An inline Water’s Acquity eλ photodiode array (PDA) detector collected spectra across a wavelength range of 190 – 340 nm. High resolution tandem mass spectrometry (MS/MS) was performed on a Q Exactive Plus mass spectrometer (Thermo Fisher Scientific, Waltham, MA) in the positive ion mode with a heated electrospray ionization source (HESI) with heater temperature at 425 °C, capillary temperature 275 °C, and spray voltage at 3.5 kV. Sheath and auxiliary gas flows were set to arbitrary values of 50 and 13, respectively. Mass spectra were collected using two scan events. The first scan was a full scan between 100 – 700 m/z at a resolution of 70,000. The second scan employed an inclusion list of the calculated m/z of all known RNA modifications as reported in the Modomics database ^43^, with a 1.0 m/z isolation window. A two-step higher-energy collisional dissociation (HCD) fragmentation of the top 5 most abundant precursor ions was employed at normalized collision energies (NCE) of 20 and 110 at 17,500 resolutions. The goal was to achieve the detection of at least two corresponding fragment ions. Before each experiment, mass calibration was performed using a canonical ribonucleoside standard mixture (3.25 µM) with an error of <1 ppm and retention time accuracy ± 0.1 min.

Each RNA sample was analyzed in triplicate, and blanks were used between repeated measurements. If multiple RNA samples were analyzed in the same experiment, the order of the repeated measurements was randomized. For each biological replicate, we also analyzed in triplicate. The structures of each RNA modification were derived from Modomics ^43^.

### 2.11 Identification of enzymatic modulators of RNA modification

To investigate the molecular basis of RNA modification changes, RNA-seq data obtained from MPLs-treated HAECs were used to quantify the expression of known RNA modification-associated enzymes, including writers (methyltransferases), readers (binding proteins), and erasers (demethylases). Differentially expressed enzymes (DEEs) were determined based on (p < 0.05) using three biological replicates per group.

### 2.12. Metabolite extraction, quantification, and LC-MS analysis

Metabolites were extracted from HAECs treated with MPLs as previously demonstrated ^44, 45^. Briefly, after incubation, cells in the 6-well plate were rinsed with 400 µl of PBS three times, then incubated with 300 µl of a 2:1 acetonitrile: water mixture (LC-MS grade) for 10 minutes on ice. Cells were then scraped, collected in Eppendorf tubes, and sonicated on ice 5 times (2-minute pulse) with a 5-second rest between each pulse. After sonication, centrifugation (16,100 × g at 4°C for 10 min) was performed to precipitate proteins. 150 µL of the resulting supernatant was transferred to autosampler vials.

Biological replicates of cellular extracts were reconstituted in acetonitrile–water followed by LCMS analysis using a Thermo Fisher Q Exactive Plus mass spectrometer (Thermo Fisher Scientific, Waltham, MA) coupled to a Waters ACQUITY ultraperformance liquid chromatography (UPLC) system (Waters Corporation, Milford, MA). A 5 µL injection volume was used for each sample, separated on a Waters ACQUITY UPLC BEH C18 column (138 Å, 1.7 µm, 2.1 × 50 mm) maintained at 40°C.

Chromatographic separation was achieved using a multi-step gradient program. The column was equilibrated at 10% B and held isocratically from 0.00 to 0.50 min. From 0.50 to 8.50 min, the organic phase was increased linearly from 10% to 100% B. This was followed by a column wash step at 100% B from 8.50 to 9.50 min. The mobile phase was then returned to initial conditions (10% B) at 9.50 min and maintained until 11.00 min to allow for column re-equilibration prior to the next injection.

After exiting the column and passing through the UV detector, the eluate was introduced into the mass spectrometer via a heated electrospray ionization (HESI) source. Data acquisition was performed in both positive and negative ion modes using full-scan MS with data-dependent MS/MS. Full-scan spectra were acquired at a resolving power of 35,000 across an m/z range of 125–1500, with an automatic gain control (AGC) target of 1×10⁶ and a maximum injection time of 100 ms.

For data-dependent acquisition, the three most abundant precursor ions in each full scan were selected for fragmentation. MS/MS spectra were collected at a resolution of 17,500 using a 1.0 m/z isolation window, an AGC target of 1×10⁵, and a maximum injection time of 50 ms. Fragmentation was performed using stepped normalized collision energies of 25, 35, and 45 eV. Additional DDA parameters included a minimum AGC target of 8×10³ for MS/MS triggering, an intensity threshold of 1.6×10⁵, dynamic exclusion set to 5.0 s, and isotope exclusion enabled ^46^.

### 2.13. LC–MS Data Processing and Feature Annotation

Raw LC–MS data files were first converted to the mzML format and subsequently processed in MZmine (version 6.3) for feature extraction. A series of modules in MZmine was applied to generate a comprehensive feature list that included feature ID, mass-to-charge ratio (m/z), retention time (RT), and peak area for each detected ion across all samples. MS/MS spectra were used for putative metabolite identification: level 2 annotation was performed by matching against the Global Natural Product Social Molecular Networking (GNPS) library, and level 3 annotation utilized the LOTUS Natural Products Database. Detailed parameters for each MZmine module are provided in **Table S2**.

To improve data quality, several filtering criteria were applied to the extracted LC–MS features. First, features with a relative standard deviation (RSD) ≤ 30% across all blank and sample runs were excluded to eliminate systematic background signals. Second, features with average signal intensity in blank injections ≥85% of the average sample signal were removed to reduce contributions from solvent or instrument-related noise. Finally, features with intra-sample variability greater than 35% (based on peak-area RSD) were discarded to retain only reproducible metabolic signals.

Following analysis with Mzmine, a total of 3132 features detected were uploaded to MetaboAnalyst as a peak intensity table for downstream preprocessing and statistical analysis. For preprocessing, feature filtering was performed using 20% RSD cutoff and 40% low-variance filter using the interquartile range (IQR). Missing values were imputed using Left-censored data estimation, with the limit of detection (LoD) set to 1/5 of the minimum positive value. Data was normalized by sum, followed by log2 transformation and auto-scaling (mean-centered and divided by the standard deviation).

Following preprocessing, differential metabolites (DMs) were identified using a two-sample t-test with unequal variances, with significance set at an FDR < 0.05. A heatmap of the top 50 metabolites was generated from normalized data using Euclidean distance and Ward linkage. We then performed PLS-DA analysis to identify important discriminating features, using VIP scores to report the top 50 features based on component 1. A volcano plot, integrating an FDR < 0.05 and a fold-change threshold of 2.0, was generated using MetaboAnalyst. For annotation, we used Sirius, and only 231 features from the T-test results were annotated.

All experiments were conducted in triplicate, and results are presented as mean ± standard error of the mean (SEM). Graphs were made using GraphPad Prism, and data were analyzed using a t-test. Statistical significance between MPLs and Ctrl group was determined at p < 0.05 (*).

## 3. Results

### 3.1. Characterization of Microplastics

Multiple techniques were used to confirm the morphology, chemical structure, and fluorescence properties of MPLs **(Fig.1a-e)**. Scanning Electron Microscopy (SEM) analysis revealed that the non-fluorescent MPLs were uniformly spherical in shape, with a mean diameter of approximately 80 nm **(Fig.1a)**. The particles exhibited smooth surfaces and minimal aggregation, indicating good monodispersity and colloidal stability under the imaging conditions.

To observe the surface chemical composition of the MPLs, Fourier Transform Infrared (FTIR) spectroscopy was performed **(Fig. 1b)**. The FTIR spectrum showed characteristic absorption bands consistent with polystyrene. Prominent peaks were observed around 3400-3300 cm⁻¹ (Primary Amine N–H stretching), 3100-3000 cm⁻¹ (Alkene C-H stretching), 1635 cm⁻¹ (C=O Alkene, Monosubstituted), 1440-1395 cm⁻¹ (O-H Carboxylic Acid, Bending), 730-665 cm⁻¹ (C=C, Alkene, Bending), 690-515 cm⁻¹, and (C-Br, Halo compound, Stretching) confirming the polystyrene backbone structure of the MPLs.

**Fig. 1.**
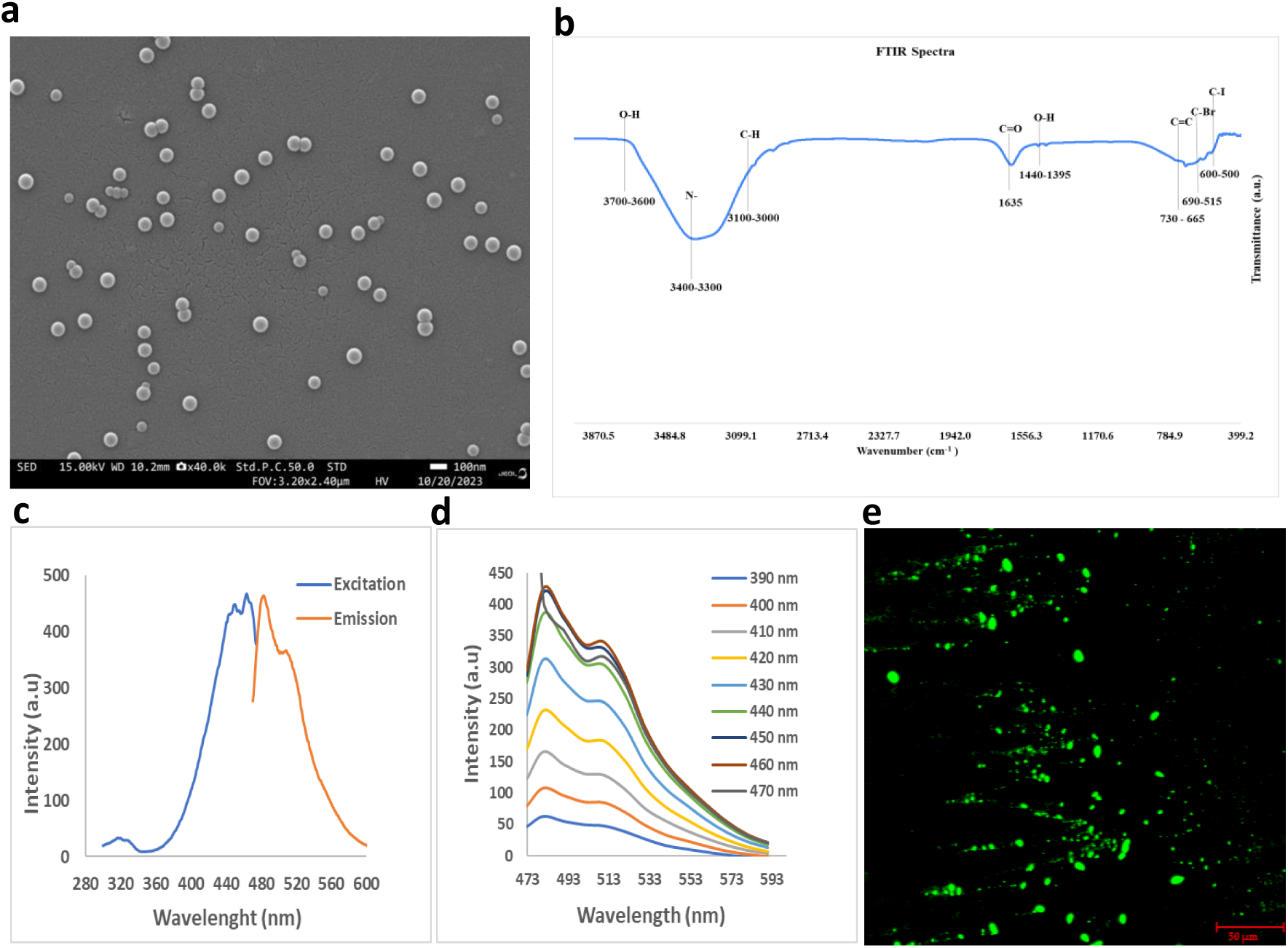
MPLs characterization: (a) Surface Morphology of MPLs viewed under Scanning Electron Microscope (SEM), highlighting the circular shape, size (80 nm), and smooth surface of MPLs, offering insights into their structural features at sub-micron resolution. (b) Fourier Transform Infrared Spectroscopy (FTIR) spectrum of MPLs. The characteristic absorption peaks shown in the spectrum correspond to functional groups found in polystyrene, confirming the integrity and chemical composition of MPLs. The prominent bands around cm⁻¹ (aromatic C–H stretching), cm⁻¹ (aromatic C=C stretching), and cm⁻¹ (C–H bending) corroborate the polystyrene structure. (c) Excitation (463.03 nm) and Emission (481 nm) peak of fluorescent MPLs (d) Emission overlay showing an emission peak of ∼481 nm with excitation wavelength of 472-592 nm of Fluorescent MPLs. (e) GFP filter image (40x) of 80 nm MPLs with Keyence BZ-X710 All-in-One Fluorescence Microscope.

The fluorescent properties of fluorescent MPLs were investigated using fluorescence spectroscopy. As shown in **Fig. 1c**, the excitation and emission spectra exhibit distinct peaks at 463.03 nm and 481 nm, respectively, indicating effective incorporation of the fluorophore into the polymer matrix. Further spectral analysis **(Fig. 1d)** revealed a strong, consistent emission peak centered at 481 nm across a range of excitation wavelengths (390–470 nm), suggesting excitation-independent emission behavior. This valuable characteristic will be employed in imaging.

To confirm the real-time fluorescence of our particles, we used Fluorescence microscopy. The GFP-filter image **(Fig. 1e)** clearly shows fluorescent MPLs as distinct green puncta, suggesting that these particles can be detected at the cell surface and may help with MPLs internalization and intracellular localization.

### 3.2. Uptake, cell viability and subcellular localization of microplastics in HAECs

To investigate cellular uptake of fluorescently labeled MPLs, HAECs were treated with 0, 10, 20, and 120 µg/mL MPLs for 24 h, and cellular uptake was quantified using a Bio-Tek® Synergy 2™ microplate reader. The uptake of MPLs by HAECs was significantly enhanced with increasing MPLs concentrations as compared to the control **(Fig. 2 a1)**. Furthermore, flow cytometry analysis supported these findings, showing a progressive rightward shift in fluorescence intensity peaks in the histogram **(Fig. 2 a2)** and a corresponding increase in the percentage of GFP-positive cells **(Fig. 2 a3)** with increasing MPLs concentration. At the highest concentration (120 µg/mL), over 80% of HAECs exhibited internalized fluorescence, confirming efficient uptake. These data collectively indicate that MPLs are readily internalized by HAECs in a dose-dependent manner. Following MPLs internalization, MTT reduction assay showed that treatment with 10, and 120 µg/mL MPLs did not significantly affect cell viability as compared to the control group **(Fig. 2 a4).**

**Fig. 2.**
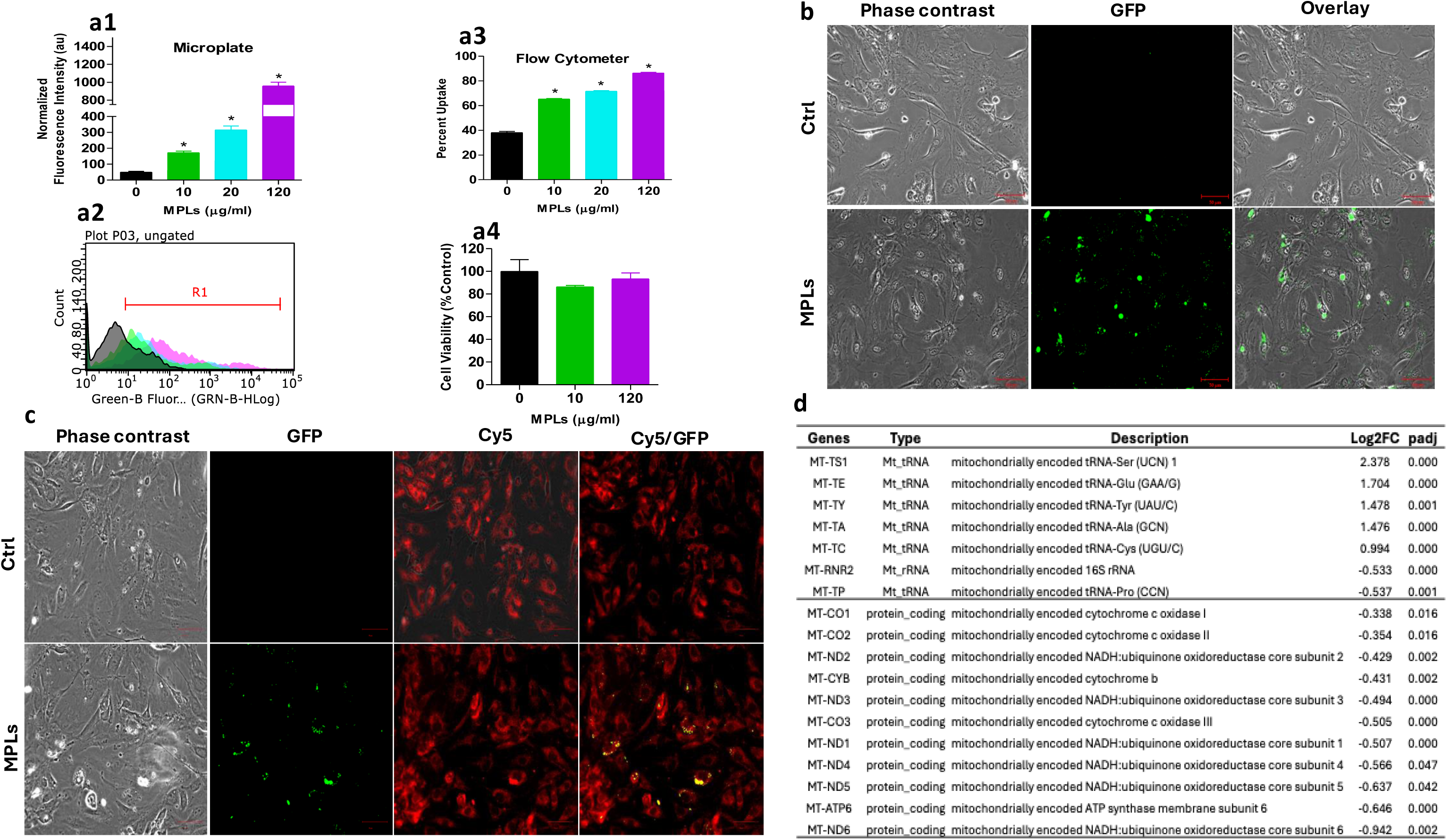
Uptake, cell viability, cellular localization of MPLs, and changes in mitochondrial genes. (a1-a3) Concentration-dependent uptake of MPLs. HAECs were treated with various concentrations (0, 10, 20, 120 µg/mL) of 80 nm MPLs for 24 h. “0” represents the vehicle-control group containing sodium azide at a final concentration of 0.00005% (w/v), without microplastics. The Bio-Tek® Synergy 2™ plate reader (a1) and flow cytometry (a2-a3) were used to quantify the uptake of MPLs. Data are expressed as mean ± SEM, n = 3. *, p < 0.05 vs. control. (a4) Cell viability following exposure to MPLs, assessed by the MTT reduction assay. (b) HAECs were treated with 0, 10 µg/mL of 80 nm MPLs for 24 h. The Keyence BZ-X710 All-in-One Fluorescence Microscope was used to visualize the localization of MPLs in HAEC cells. Representative (n=3 and field of view = 9 per condition) phase-contrast, GFP (green), and overlay images indicate that MPLs are taken up by HAEC cells. (c) Representative images showing MPLs mitochondrial localization in HAECs: Endothelial cells were pre-incubated with 0.5μM of MitoTracker™ Red CMXRos for 30 min and then treated with and without MPLs (Size: 80 nm, 10 µg/mL) for 24 h. The Keyence BZ-X710 All-in-One Fluorescence Microscope was used to visualize the interaction of MPLs with mitochondria in HAEC cells. Representative (n=3 and field of view = 9 per condition) phase-contrast (PC), GFP (green), Cy5 (red), and overlay (PC/GFP, Cy5/GFP) images indicate that MPLs interact with endothelial mitochondria. (d) Transcriptomics data of MPLs-treated HAECs show altered expression of mitochondrial protein-coding genes and mitochondrially encoded tRNA genes as compared to the control. Values are shown as Log2FC and adjusted p values (padj < 0.05).

To further confirm the results of the microplate reader and flow cytometer, HAECs treated with fluorescently labeled MPLs for 24 h were imaged using fluorescence microscopy. Phase contrast, GFP, and overlay imaging **(Fig. 2b)** revealed a dose-dependent uptake of MPLs, thereby corroborating the results from the microplate reader and flow cytometer. Cells exposed to 10 µg/mL of MPLs (**Fig. 2b**, Phase contrast and GFP and overlay) demonstrated a substantial accumulation of green fluorescence throughout the cytoplasm, whereas untreated control cells **(Fig. 2b)**, Phase contrast, GFP and overlay imaging) showed no detectable fluorescence signal. Overlay image of MPLs treated Phase contrast and GFP image **(Fig. 2b)** confirmed that MPLs were internalized rather than adhering to the extracellular surface.

To explore the subcellular fate of internalized MPLs, we evaluated their co-localization with mitochondria using Mitotracker™ Red CMXRos labeling. Following 24-hour exposure to MPLs (10 µg/mL), cells were imaged for GFP (green, MPLs) and Cy5 (red, mitochondria) fluorescence **(Fig. 2c)**. In untreated cells (**Fig. 2c**; 0 µg/mL), no green fluorescence was detected. However, in MPLs-treated cells (**Fig. 2c**; 10 µg/mL), the internalized MPLs exhibited substantial co-localization with mitochondrial structures, as evidenced by green puncta in the merged Cy5/GFP channels (**Fig. 2c**; 10 µg/mL). Multiple mitochondrially encoded tRNAs were among the most strongly induced transcripts (**Fig. 2d**), including MT-TS1 (log2FC = 2.38, padj < 0.001), MT-TE (1.70, <0.001), MT-TY (1.48, =0.001), MT-TA (1.48, <0.001), and MT-TC (0.99, <0.001). In contrast, the majority of protein-coding genes that form the electron transport chain were significantly reduced. Components of complex I showed broad suppression, such as MT-ND1, MT-ND3, MT-ND4, MT-ND5, and MT-ND6 (**Fig. 2d**). Similar decreases were observed for complex IV genes MT-CO1, MT-CO2, and MT-CO3, as well as MT-CYB of complex III and the ATP synthase subunit MT-ATP6, indicating that MPLs exposure is associated with repression of oxidative phosphorylation in HAECs. These findings suggest that MPLs are not only efficiently taken up by HAECs but may also associate with or localize near mitochondrial compartments, implicating potential organelle-specific interactions and functions.

MPLs localization with lysosome was studied by staining lysosome with LysoTracker Deep Red dye **(Fig. 3)**. Confocal live-cell imaging (transmitted-light [TD], GFP, LysoTracker Deep Red, and merged channels) of the Ctrl group showed low background signal in both GFP and Deep Red channels, with no evident punctate structure in merged images. In the MPLs-only group, a clear GFP-positive signal was observed throughout the cytoplasm, and no Deep Red signal was observed, showing that the green signal reflected MPLs-associated fluorescence and not the lysosomal staining. Whereas, in the lysosome-only group, we observed LysoTracker Deep Red prominent punctate/perinuclear vesicular staining, which was consistent with lysosomal compartments, and no GFP signals were observed. Importantly, in the MPLs+lysosome group, we observed clear, strong signals in both channels, and the merged images showed clear spatial overlap between GFP-positive MPLs puncta and LysoTracker-positive vesicles that appeared as co-localized puncta within the lysosome-rich region. We also observed clear co-localization in perinuclear and intracellular vesicular compartments, consistent with endolysosomal trafficking and lysosomal accumulation of MPLs after 24 h of exposure.

**Fig. 3.**
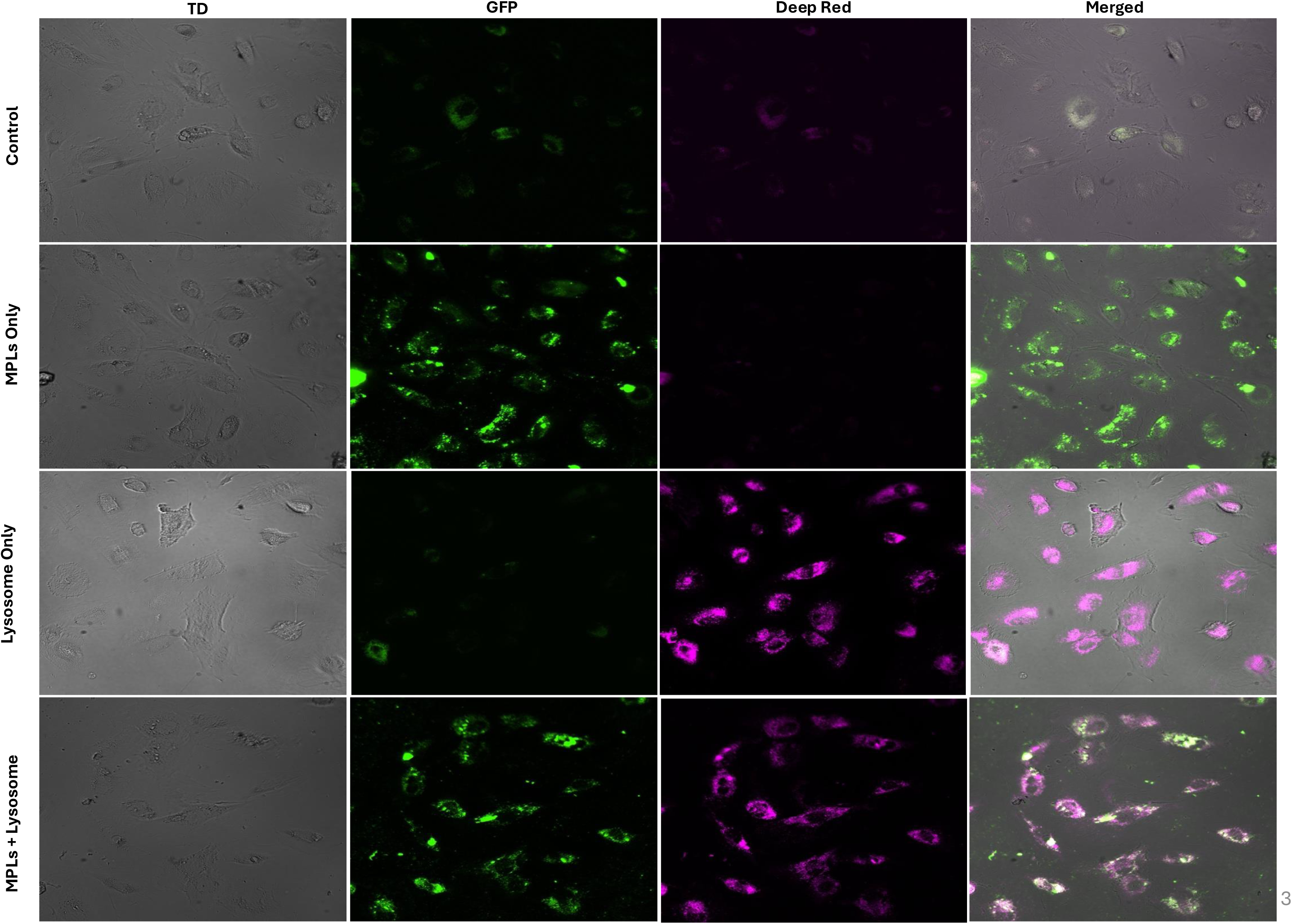
Microplastics show intracellular co-localization with lysosomes in HAECs: Representative live-cell confocal images of HAECs showing transmitted light (TD), microplastics (MPLs; GFP channel, green), lysosomes (LysoTracker® Deep Red, magenta), and merged overlays. Cells were imaged under four conditions: Control, MPLs only (24 h), LysoTracker only (100 nM, last 2 h), and MPLs + LysoTracker (MPLs for 24 h with LysoTracker added for the final 2 h). In the MPLs + LysoTracker condition, merged images show overlap of GFP-positive MPL signal with LysoTracker-positive puncta, consistent with MPLs localization within lysosome-associated compartments. Images were acquired using a Nikon AXR resonant confocal microscope (40× air objective) under live-cell conditions (37°C, 5% CO₂).

### 3.3. Identification of the cellular uptake pathway of MPLs

Elucidating the possible uptake pathways of exogenous substances (such as MPLs) is crucial for revealing their physiological effects. Previous studies have shown that endocytosis is one of the key pathways for nanoparticle uptake ^47^. Based on the concentration determined in our previous study, human aortic endothelial cells (HAECs) were treated with fluorescently labeled polystyrene MPLs (MPLs, 10 µg/mL) and then co-incubated with various ion channel blockers and endocytosis inhibitors **(Table S1)** for 30 minutes to investigate the potential uptake and internalization pathways of MPLs **(Fig. 4 a1-j)**.

**Fig. 4.**
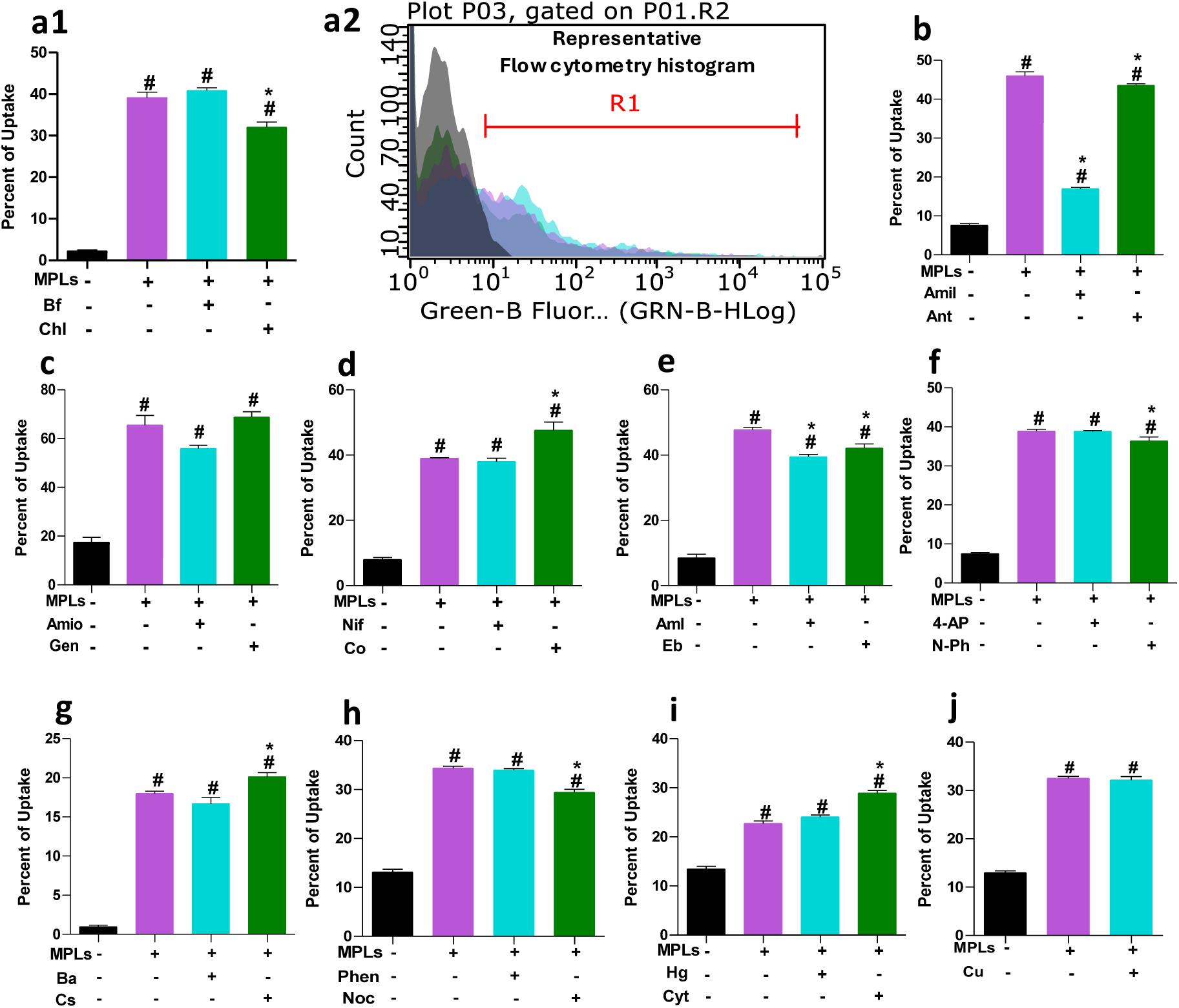
Effect of various ion channel blockers and endocytic inhibitors on uptake of MPLs in HAECs. HAECs were treated with ion channel blockers and endocytic inhibitors for 30 minutes, then incubated with MPLs (80 nm, 10 μg/mL) for 24 h. Guava Soft Flow cytometer was used to quantify the uptake of MPLs. (a1, b-j) bar graph showing quantified mean fluorescent percent intensity values, and (a2) shows the representative flow Cytometry histogram. Data are expressed as mean ± SEM (n = 3. # p < 0.05 vs Ctrl, * p < 0.05 vs MPLs).

Compared to the control group, HAECs exposed to fluorescent MPLs (10 µg/mL) alone and detected by Guava® easyCyte™ flow cytometry (Millipore™) showed significant MPLs internalization (p < 0.05) **(Fig. 4 a1)**. However, co-treatment of HAECs with chlorpromazine (Chl), a known inhibitor of clathrin-mediated endocytosis, 30 minutes before MPLs treatment, significantly reduced microplastic uptake (*p < 0.05); while treatment with bafilomycin A1 (Bf), a specific vacuolar H⁺-ATPase (V-ATPase) inhibitor used to inhibit endosomal and lysosomal acidification, did not cause a significant reduction in uptake **(Fig. 4 a1)**. The representative flow cytometry histogram overlay results further confirmed that, compared to the control group (gray curve), the green fluorescence channel signal of the MPLs group (pink curve) shifted to the right, indicating increased MPLs uptake by the cells **(Fig. 4a2)**. After co-treatment with Chl, the cellular fluorescence intensity decreased and the curve shifted to the left compared with MPLs alone **(Fig. 4a1)**, further illustrating its inhibitory effect on uptake.

Co-treatment with amiloride hydrochloride (Amil, an inhibitor that inhibits macropinocytosis by blocking the Na⁺/H⁺ exchange pump) significantly reduced the uptake of MPLs (* p < 0.05) as compared to the MPLs alone **(Fig. 4b; cyan curve).** Anthracene-9-carboxylic acid (Ant), a chloride channel blocker, also significantly (* p < 0.05) reduced the uptake of MPLs as compared to the MPLs alone **(Fig. 4b; green curve)**. Treatment with amiodarone hydrochloride (Amio) and genistein (Gen), the latter being a tyrosine kinase receptor inhibitor, did not significantly affect microplastic uptake **(Fig. 4c)**. Niflumic acid (Nif, 10 µM) treatment also did not alter the uptake level **(Fig. 4d)**, and cobalt chloride (Co), a non-selective inorganic calcium-channel blocker, did not significantly reduce MPLs uptake **(Fig. 4d)**. Amlodipine (Aml, a dihydropyridine L-type calcium channel blocker), and ebselen (Eb, a mammalian H⁺/K⁺-ATPase inhibitor), both significantly (* p < 0.05) reduced microplastic uptake (p < 0.05) **(Fig. 4e)** confirming the reduced uptake of MPLs in response to these inhibitors. Treatment with 4-aminopyridine (4-AP, a potassium channel blocker) resulted in uptake levels comparable to the MPLs group **(Fig. 4f)**, while N-phenylanthranilic acid (N-Ph, a chloride channel blocker) significantly (* p < 0.05) reduced MPLs uptake (p < 0.05) as compared to MPLs alone **(Fig. 4f; green curve)**. Treatment with anhydrous barium chloride (Ba) did not significantly alter the uptake rate **(Fig. 4g)**. Likewise, treatment with cesium chloride (Cs, a potassium channel blocker) did not reduce MPLs uptake (**Fig. 4g)**. treatment with phenylglyoxal (Phen, a selective inhibitor of phagocytosis) also did not significantly affect uptake **(Fig. 4h)**. However, nocodazole (Noc, a microtubule and actin disruptor) significantly (* p < 0.05) reduced uptake as compared to the MPLs alone group **(Fig. 4h; green curve).** Treatment with mercuric chloride (Hg, an hAQP1 aquaporin inhibitor) and cytochalasin A (Cyto, an actin disruptor) did not significantly reduce MPLs uptake **(Fig. 4i)**. Finally, treatment with copper sulfate (Cu, a hAQP3 aquaporin inhibitor) did not have a significant effect on uptake **(Fig. 4j)**.

In summary, the results of the inhibitor effects on microplastic uptake by HAECs indicate that chlorpromazine (Chl), amiloride (Amil), anthracene-9-carboxylic acid (Ant), amlodipine (Aml), ebselen (Eb), N-phenylanthranilic acid (N-Ph), and nocodazole (Noc) can reduce microplastic uptake, suggesting that the specific ion channels or endocytic pathways corresponding to these inhibitors play an important role in the uptake of MPLs by HAECs.

### 3.4. Transcriptomic analysis reveals enrichment of endocytic and vesicular transport pathways in MPLs HAECs

Using bulk RNA sequencing, we systematically identified key genes and their functional modules involved in cellular uptake mechanisms via differential expression and pathway enrichment analyses. Results of transcriptome analysis showed that multiple genes involved in endocytosis, actin dynamics, and vesicle transport (i.e., cytoskeleton remodeling) were significantly upregulated in the MPLs treatment group **(Fig. 5a)**. Compared to the control (0 µg/mL), the expression of genes encoding actin-binding proteins (such as *ACTBP2*, *ACTG1P9*) and Rho GTPase regulatory factors (such as *ARHGAP20*, *ARHGAP22*, *RAC1P2*, *RAC2*, *RAC3*) was significantly increased (p < 0.05) in MPLs-treated cells **(Fig. 5a)**. Simultaneously, transcripts encoding Rab family GTPases (such as *RAB6C*, *RAB6D*, *RABAC1*, *RAB32*) and ESCRT and sorting connector complex components (such as *VPS18*, *VPS25*, *SNX2*, *SNX8*) also showed a significant increase (p < 0.05, **Fig. 5a**). These changes are consistent with our previous cellular uptake experimental results (**Fig. 2 a1-c**), showing the systematic activation of transcriptional programs related to vesicle formation and intracellular transport under MPLs stimulation.

**Fig. 5.**
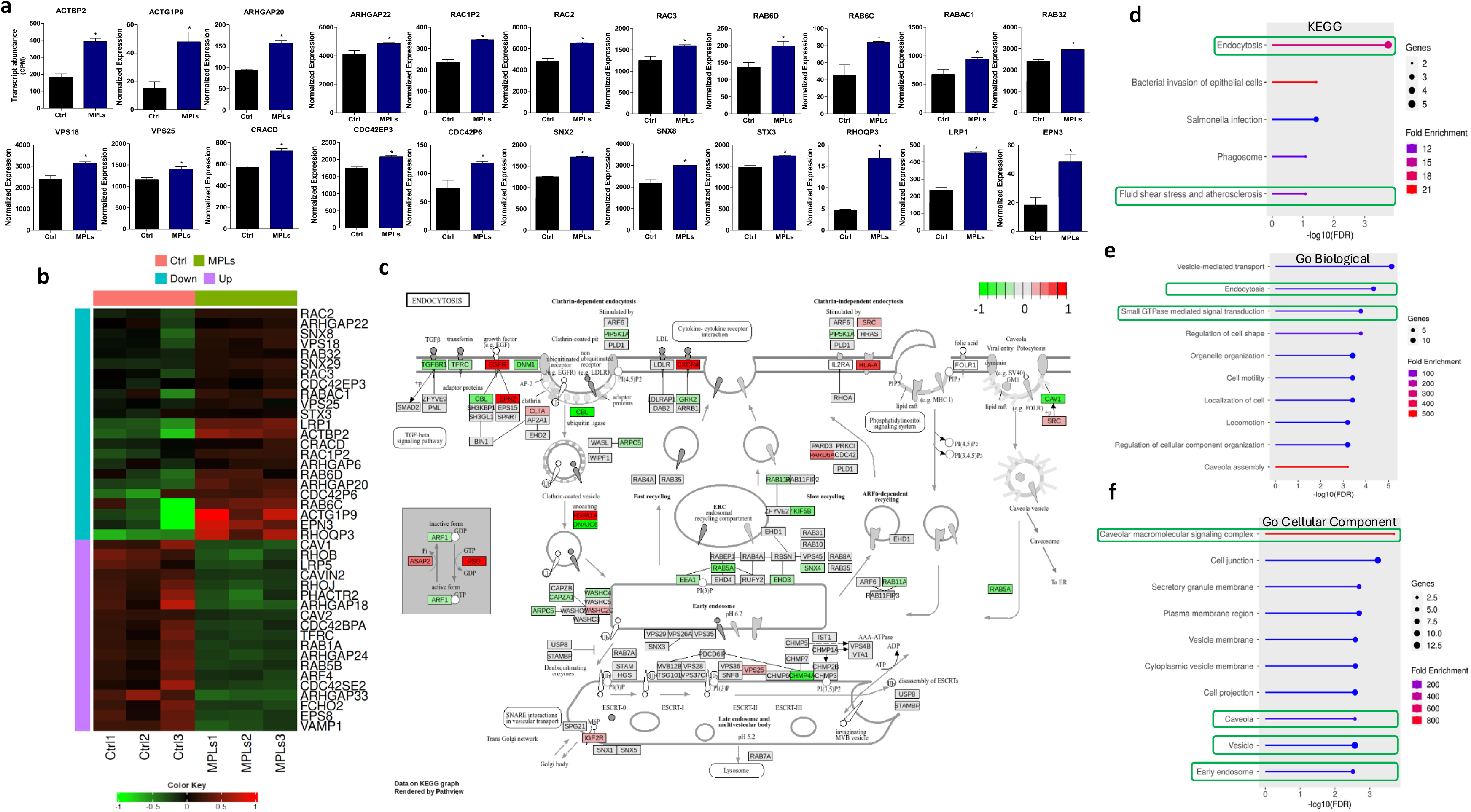
Expression profiles of genes involved in MPLs uptake: (a) Bar plot represents the relative expression of key genes involved in MPLs internalization pathways following MPLs treatment. Gene expression was quantified using RNA Sequencing. Each plot represents an individual gene involved in cellular uptake mechanisms, such as endocytosis, phagocytosis, and receptor-mediated transport. Data presented as mean ± SEM from three biological replicates. Normalized counts are shown as mean ± SEM (n = 3 biological replicates). All genes shown were significantly differentially expressed between control and MPLs-treated cells in the DESeq2 analysis. * padj < 0.05 compared to the control group. (b) Heat map of DEGs involved in uptake (c) Kegg Endocytosis pathway with significantly changed gene involved in endocytosis are highlighted (upregulated in red, downregulated in green). Lollipop plot of KEGG pathway enrichment. (d) KEGG pathway, (e) GO Biological Processes and (f) Go cellular Component enrichment highlighting the involvement of the differentially expressed genes in MPLs internalization.

The heatmap generated by hierarchical clustering further showed that MPLs HAECs exhibited a unique gene expression pattern, with widespread upregulation of genes involved in membrane remodeling, vesicle sorting, and cytoskeletal organization **(Fig. 5b)**. Among them, genes such as *EPN3*, *RHOQP3*, *CDC42P6*, and *LRP1* showed upregulation, suggesting that multiple aspects of endocytosis were activated.

At the pathway level, cellular uptake and vesicle trafficking DEGs (43) showed significant enrichment in the endocytosis pathway. Projecting the significantly altered genes onto the KEGG endocytosis pathway map revealed transcriptional involvement in both clathrin-dependent and clathrin-independent endocytosis pathways **(Fig. 5 d)**. Genes involved in key nodes such as cargo recognition (e.g., *TFRC*, *EGFR*), vesicle fission (e.g., *DNM1*, *EPS15*), early endosome transport (e.g., *RAB5A*, *EEA1*), and membrane recycling all showed an upward trend. These data collectively suggest that both classical and non-classical endocytic pathways play important roles in the transcriptional response to MPLs exposure.

Finally, gene ontology (GO) and KEGG enrichment analyses of 43 DEGs involved in cellular uptake and vesicle trafficking further confirmed endocytosis as the main internalization pathway **(Fig. 5c, d)**. KEGG analysis showed that endocytosis is the main internalization pathway (with genes from our RNA Seq data highlighted in Red and Green), and fluid shear stress-related atherosclerosis pathways were the most significantly enriched pathway categories **(Fig. 5c, d)**. GO biological process analysis revealed that terms related to vesicle-mediated transport, endocytosis, and cell morphology regulation were significantly enriched after MPLs treatment **(Fig. 5e)**. GO cellular component analysis revealed that terms related to structures such as caveolae, early endosomes, vesicle membranes, and caveolae macromolecular signaling complexes were the most highly enriched **(Fig. 5f)**. The caveolae-associated enrichment was observed at the transcriptomic level only.

In summary, this study demonstrates that HAECs undergo a series of coordinated transcriptional reprogramming after exposure to MPLs, involving the systematic activation of pathways related to endocytosis, intracellular vesicle transport, and mechanotransduction.

### 3.5. Transcriptomic Profiling Identifies Differential Gene Expression in MPLs Exposed HAECs

Principal component analysis (PCA) of global transcriptomic profiles revealed distinct separation between MPLs-treated and control human aortic endothelial cells (HAECs), with PC1 accounting for 87.84% and PC2 for 6.27% of total variance **(Fig. 6a)**. All MPLs-exposed replicates (MPLs 1–3) clustered distinctly from the control samples (Ctrl 1–3), showing a difference in gene expression following MPLs treatment. Classification of detected transcripts by gene type indicated a predominance of protein-coding genes (n = 12,565), followed by long non-coding RNAs (lncRNAs, n = 2,114) and pseudogenes (n = 1,389) **(Fig. 6b)**. Others included microRNAs (miRNAs), small nucleolar RNAs (snoRNAs), small nuclear RNAs (snRNAs), and mitochondrial RNAs. Differential expression analysis identified 1,168 significantly upregulated and 926 significantly downregulated differentially expressed genes (DEGs) in response to MPLs exposure **(Fig. 6c)**. These were defined by statistical thresholds applied to log₂ fold-change and adjusted p-values. The volcano plot **(Fig. 6d)** highlighted genes exhibiting both high magnitudes and statistical significance in expression changes. Among the most significantly upregulated genes were *MMP1, FOSB, STC1, CDCP1, HMOX1,* and *LYVE1*. Conversely, transcripts such as *CASP12, GDF7, CX3CL1,* and *PDE1A* were among the most significantly downregulated. A heatmap of the top 100 differentially expressed genes (DEGs) **(Fig. 6e)** showed clear clustering between MPLs-treated samples and the Ctrl group, demonstrating the magnitude and directionality of gene regulation and highlighting consistent transcriptional shifts across replicates. Tabulation of the top 20 upregulated and top 20 downregulated genes is presented in **Fig. 6f**. Genes such as *HSPA7, HSPA6, OASL, FOSB, CCL5, STC1,* and *MMP10* showed the highest positive log₂ fold-change values (ranging from 2.07 to 7.96), with padj < 0.01. Downregulated transcripts included *CASP12*, *GDF7*, *CX3CL1*, *KCNT2*, and *PDE1A*, exhibiting negative log₂ fold-change values and significant padj values. These findings collectively establish a comprehensive gene expression profile associated with MPLs exposure, characterized by significant modulation of genes involved in diverse cellular processes.

**Fig. 6.**
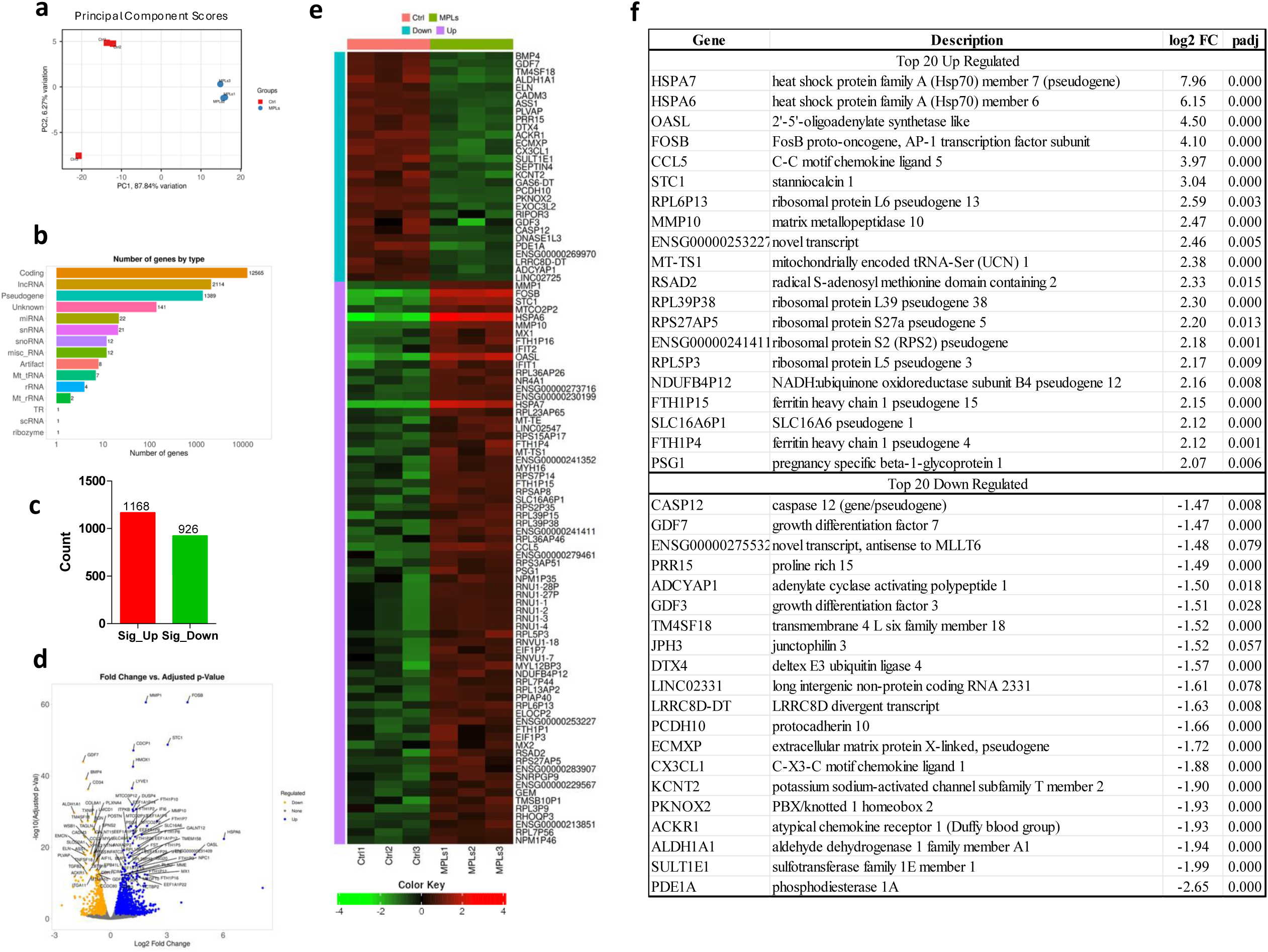
Global qualitative analysis and differentially expressed transcriptomics profile of HAECs treated with MPLs: (a) Principal component analysis (PCA) on Padj < 0.05 data reveal that PC1 accounts for 87.8% of the total variance, demonstrating a clear difference between the Control and MPLs Group. PCA also shows good clustering of replicates, demonstrating experimental reproducibility and validity, and shows a clear effect of MPLs on the transcriptomics profiles of HAEC cells. (b) The bar plot shows that 12,565 of the total genes were coding genes. (c) Bar plot of significant DEGs (padj < 0.05) demonstrates 1168 upregulated and 926 downregulated genes in response to MPLs-treated cells. (d) A volcano plot of 100 annotated DEGs shows significant up- and downregulated genes. (e) Heat Map of top 100 DEGs sorted by Fold Change (up and down regulated) showing a clear effect of MPLs on transcriptomics profile of HAECs. (f) The table of the top 20 strongly up- and downregulated DEGs, with their descriptions, shows that upregulated genes are involved in extracellular matrix remodeling, stress response, and angiogenic factors. The table of the top 20 downregulated DEGs, with descriptions, shows their involvement in various processes of vesicle trafficking.

Transcriptomics and functional analysis reveal activation of the inflammatory signaling pathway via NF-κB following exposure to MPLs. Gene Ontology (GO) enrichment analysis of upregulated transcripts revealed overrepresentation of biological processes related to host defense and external stimuli response **(Fig. 7a)**. The most enriched terms included response to external stimulus, response to viruses, and Negative regulation of viral genome replication. Additional categories, such as Immune system process, Immune response, and response to biotic stimulus, were also enriched. The number of contributing genes and corresponding fold enrichment values are indicated by point size and color, respectively, with −log₁₀(FDR) used for ranking. Gene Set Enrichment Analysis (GSEA) based on the MSigDB Hallmark collection identified HALLMARK_ TNF-α_SIGNALING_VIA_NFKB as the top positively enriched pathway in MPLs-treated HAECs, with a normalized enrichment score (NES) of 1.6 and padj value of 0.003 **(Fig. 7b and 7c)**. Additional enriched Hallmark gene sets included INTERFERON_ALPHA_RESPONSE, INTERFERON_GAMMA_RESPONSE, IL6_JAK_STAT3_SIGNALING, and P53_PATHWAY. Positively enriched pathways are indicated by positive NES values. Downregulated gene sets, such as EPITHELIAL_MESENCHYMAL_TRANSITION and E2F_TARGETS, are represented by negative NES values **(Fig. 7b)**. The enrichment plot for TNF-α signaling via NF-κB **(Fig. 7c)** shows a peak enrichment score corresponding to the leading-edge genes contributing to the signal. These include *FOSB*, *IFIT2*, *NR4A1*, *SQSTM1*, *TNFSF9*, *FOSL1*, *JUN*, *EGR1*, *IL15RA*, *CD44*, and *KLF4*. GO analysis of downregulated transcripts identified enrichment in biological processes associated with cell structure and morphogenesis **(Fig. 7d)**. Cell adhesion, Anatomical structure morphogenesis, Cell migration, and Tissue development were among the top categories. These terms had statistically significant −log₁₀(FDR) values and were associated with a high number of downregulated genes, as indicated by dot size and color. Reactome pathway analysis **(Fig. 7e)** demonstrated differential regulation of multiple cellular processes. Upregulated pathways (highlighted in green) involved Interferon signaling, Antiviral mechanisms by IFN-stimulated genes, and Interferon alpha/beta signaling. Downregulated pathways (highlighted in red) included RNA Polymerase II Transcription, Extracellular matrix organization, Nucleosome assembly, and Histone methylation. Luciferase reporter assay for NF-κB activity **(Fig. 7f)** revealed a significant (p < 0.05) increase in relative luminescence units (RLU) in HAECs treated with MPLs (120 µg/mL) compared to the control. This result confirms transcriptional activation of NF-κB in response to MPLs exposure.

**Fig. 7.**
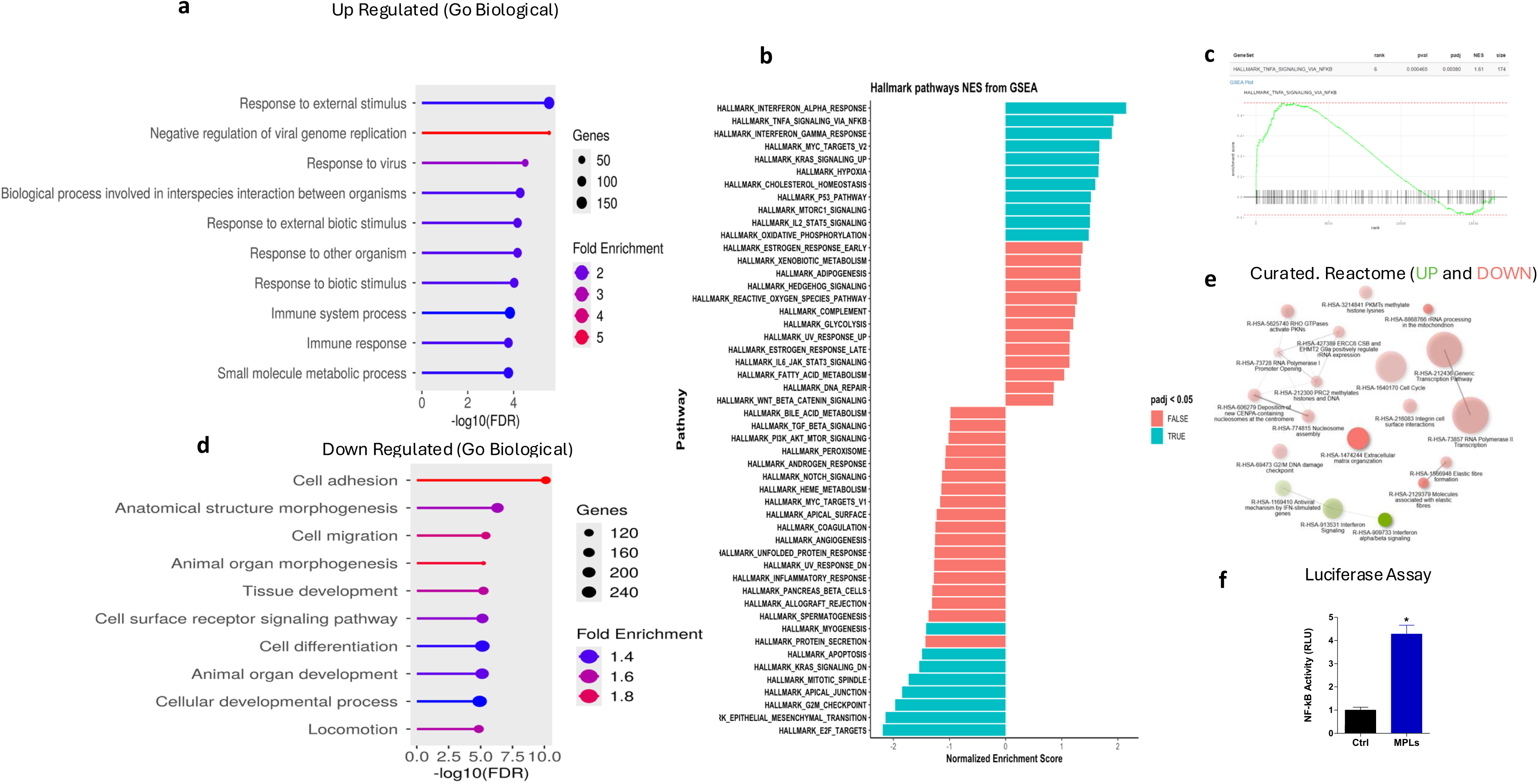
The Modulation of the Inflammatory pathways by MPLs in HAECs. (a) A lollipop plot of upregulated biological processes in response to MPLs treatment shows many terms clustering around positive regulation of gene expression, defense against viruses, and immune response, demonstrating an inflammatory signature linked to atherosclerosis. Circle size indicates the number of genes involved, and color indicates fold enrichments. (b) Gene set enrichment analysis (GSEA) ranking Hallmark pathways by normalized enrichment score (NES) shows that TNF-α signaling via NFkB is the most dominant pathway, positively enriched (p < 0.05). Several other interferons and an inflammatory response signature are also enriched. (c) GSEA was performed using MSigDB Hallmark gene collection (h.all, human), demonstrating that HALLMARK_TNFA_SIGNALING_VIA_NFKB gene set was significantly enriched in the MPLs condition (padj = 0.003; NES = 1.6; gene set size = 174). The enrichment line (green) and the positions of genes from the gene set along the ranked list of all genes are shown. Vertical lines represent individual genes, and the peak of the green line shows maximum enrichment. The key genes contributing to the enrichment signal include *FOSB*, *IFIT2*, *NR4A1*, *SQSTM1*, *TNFSF9*, *FOSL1*, *JUN*, *EGR1*, *IL15RA*, *CD44*, and *KLF4*, which are responsible for pro-inflammatory response. (d) Lollipop of downregulated GO Biological pathways reveals suppression of transcriptional and RNA metabolic programs. MPLs halting these machineries are signatures of chronic vascular inflammation. (e) Human Reactome Up- and Downregulation pathways. (f) Luciferase reporter NFKB activity demonstrates increased NFKB activity, validating the role of MPLs in activating the pro-inflammatory NFKB pathway.

MPLs exposure caused a strong pro-atherogenic response in HAECs demonstrated by upregulation of genes that are involved in processes that could lead to atherosclerosis such as inflammation and immune response (*CCL5*, *IL33*), leukocyte-adhesion/immune regulation genes (*ICAM2*, *CD44*, *CD276*, *CXCL16*/*CXCR4*) and others **(Fig. 8a)**. In parallel, MPLs suppressed key protective pathway genes such as antioxidant and stress-adaptation programs (*NFE2L1*, *SOD2*, *GPX3*, *HSPA2*/*HSPA5*/*HSP90B1*, *CREB1*) and lipid/cholesterol homeostasis regulators (*ABCA6*, *INSIG2*) **(Fig. 8b)**. Furthermore, MPLs significantly upregulated mitochondrially encoded tRNAs such as *MT-TS1*, *MT-TE*, *MT-TY*, *MT-TA*, *MT-TC*, while MT-RNR2 and MT-TP were downregulated. However, most mitochondrially encoded electron transport chain/oxidative phosphorylation genes were reduced, including *MT-CO1*/*CO2*/*CO3*, *MT-ND1*/*2*/*3*/*4*/*5*/*6*, *MT-CYB*, and *MT-ATP6*, demonstrating suppression of mitochondrial respiratory gene expression following MPLs Treatment **(Fig. 2d)**.

**Fig 8.**
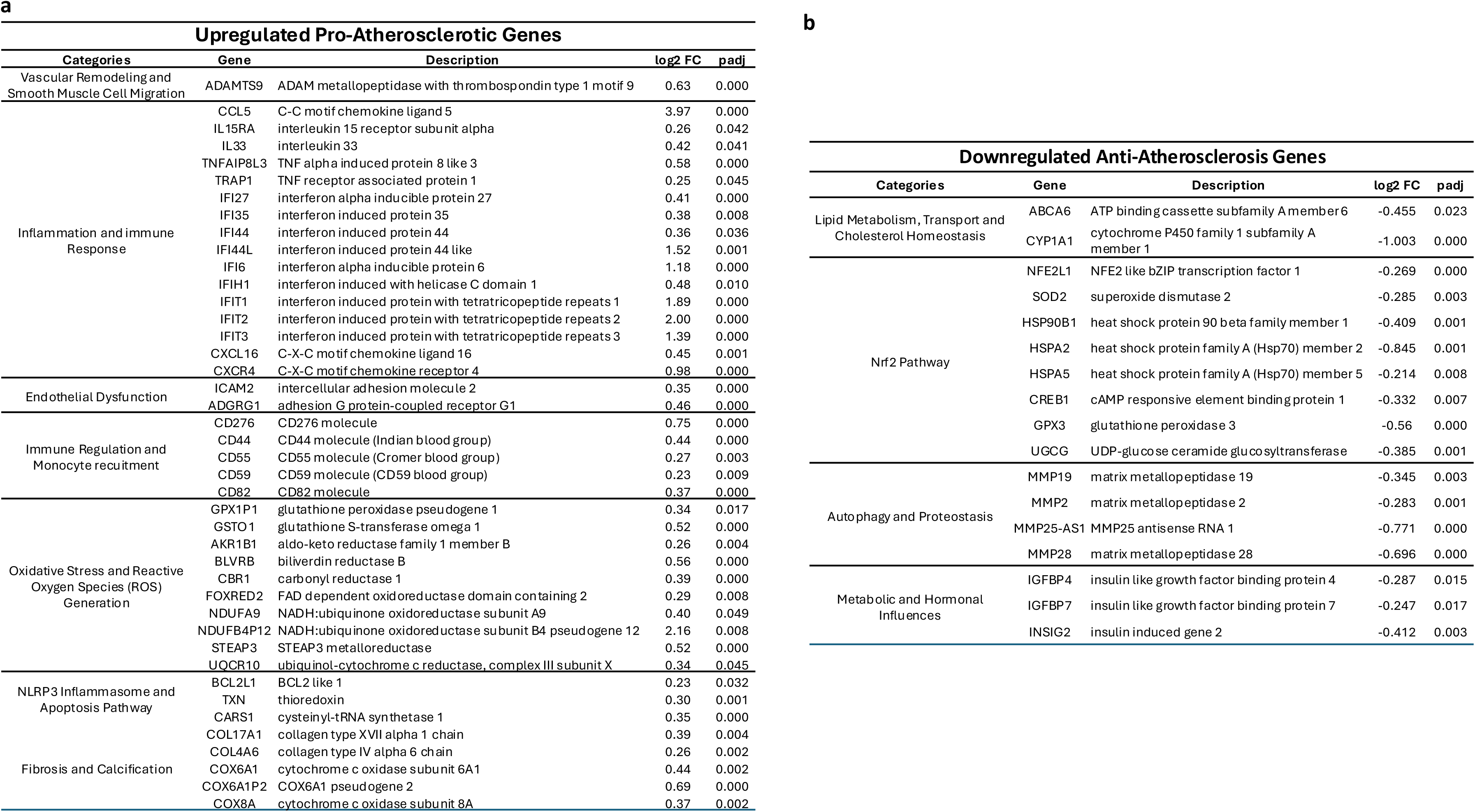
MPLs induced DEGs of interest reprograms HAEC cells to a pro atherogenic condition: (a) Up regulation (padj < 0.05) of pro atherosclerotic genes in response to MPLs treatments are clearly involved in process which could lead to atherosclerosis such as vascular remodeling and smooth muscle cell migration, inflammation and immune response, endothelial dysfunction, immune regulation and monocyte recruitment, oxidative stress and reactive oxygen species (ROS) generation, NLRP3 Inflammasome and apoptosis pathway and fibrosis and calcification. (b) Downregulation (padj < 0.05) of anti-atherosclerosis genes demonstrates that, in response to MPLs, genes and processes that protect against atherosclerosis are downregulated, including lipid metabolism, transport, and cholesterol homeostasis; the Nrf2 pathway; autophagy and proteostasis; and metabolic and hormonal Influences. Both tables clearly demonstrate that, in response to MPLs, genes are activated that could activate pathways and processes leading to atherosclerosis, and genes and pathways are suppressed that are anti-inflammatory, pointing to atherosclerotic reprogramming of HAEC cells in response to MPLs.

Prize-Collecting Steiner Forest (PCSF) network reconstruction through xOmics Shiny server (https://xomicsshiny.bxgenomics.com/) reveals multimodal inflammatory and vascular remodeling signatures in MPLs treated HAECs (**Fig. 9)**. Using PCSF (fold change ≥ 1.8, padj < 0.05), transcriptomic data from MPLs treated HAECs were mapped to identify functionally coherent gene clusters. The resulting network contained 22 distinct signaling modules, among which seven clusters exhibited strong relevance to cardiovascular inflammation and remodeling. Notably, the orange cluster (*CXCR4, JAK2, HIF1A*) highlighted cytokine- and hypoxia-mediated TNF-α and JAK-STAT signaling, while the dark green cluster (*IFI6, IRF9, NFKB1*) captured type I interferon- and NF-κB-driven immune responses. Additional clusters were enriched for TGF-β/ECM remodeling (green), interferon-stimulated genes and adhesion molecules (yellow), chemokine-mediated immune recruitment (pink), MAPK/NF-κB stress signaling (dark red), and BMP-driven vascular patterning (blue). Terminal nodes represented differentially expressed genes, and Steiner nodes indicated intermediate signaling components. Together, the network reveals coordinated activation of pro-inflammatory, immunomodulatory, and vascular remodeling pathways in HAECs following MPLs exposure.

**Fig. 9.**
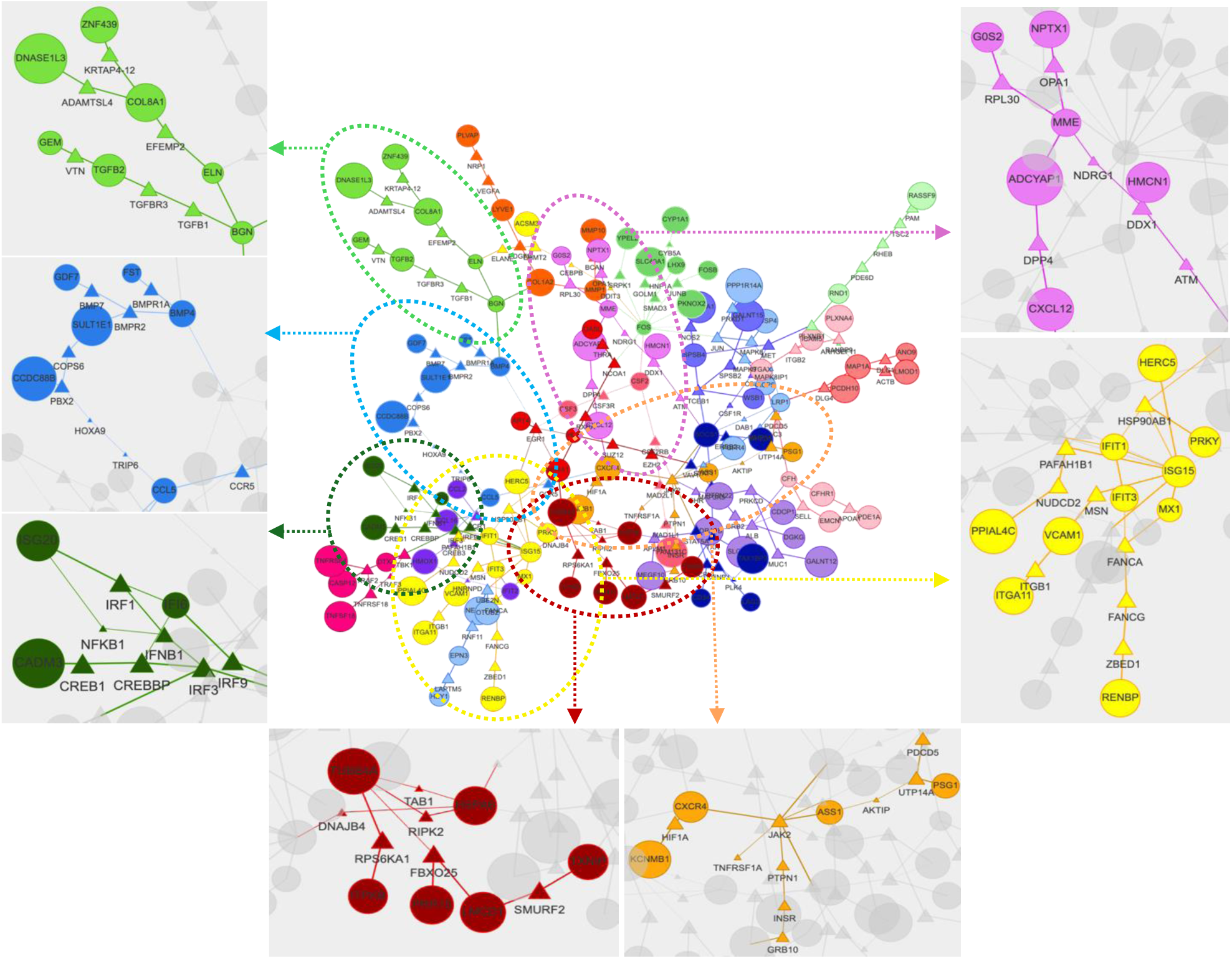
Prize-Collecting Steiner Forest (PCSF) network reconstruction demonstrates the role of MPLs in HAEC signaling clusters linked to inflammation, immune recruitment, and vascular remodeling. MPLs treated HAECs were analyzed using the PCSF network reconstruction with KEGG annotations (Fold change: 1.8 and padj < 0.05), revealing functionally distinct gene clusters (color-coded) and their roles in conditions (such as inflammation, stress, vascular remodeling) culminating in atherosclerosis (cardiovascular diseases). Terminal Nodes (Circles) represent DEGs, and Steiner Nodes (Triangles) are connecting genes. Among 22 clusters, seven clusters demonstrate strong functional relevance to cardiovascular inflammation and atherosclerosis: 1) Orange Cluster: Genes such as *CXCR4, JAK2, TNFRSF1A, ASS1, and HIF1A in this cluster are centered around cytokine receptor and hypoxia signaling, showing TNF and JAK-STAT pathways activation leading to inflammation, vascular remodeling, and endothelial dysfunction in cardiovascular diseases and atherosclerosis. Dark Green Cluster: Genes such as IFI6, IFNB1, IRF1, IRF3, IRF9, NFKB1, and ISG20* in this cluster are involved in type 1 interferon and NFKB immune responses, which are activated in response to endothelial dysfunction, immune cell infiltration, and plaque destabilization. Green Cluster: Genes *TGFB1, ELN, COL8A1, BGN, EFEMP2, and TGFBR3* in the cluster are involved in TGF-β signaling and extracellular matrix (ECM) remodeling required for vascular stiffening and intimal thickening, signatures of atherosclerosis. Yellow Cluster: Genes such as *ISG15, IFIT1, IFIT3, VCAM1, ITGA11, ITGB1, and MX1* in this cluster are involved in processes such as interferon-stimulated genes and cell adhesion molecules, suggesting vascular inflammation. Pink Cluster: Genes such as *CXCL12*, *DPP4*, *MME*, *ADCYAP1*, and *NPTX1* are enriched in chemokine-mediated immune recruitment, suggesting MPLs, which promote immune cell attachment and inflammation. Dark Red Cluster: *RIPK2*, *HSPA6*, *TAB1*, *TUBB4A*, and *PRR15* genes in this cluster are involved in inflammatory stress response and apoptosis, including MAPK/NF-κB signaling effectors, indicating cell stress and remodeling. Blue Cluster: Genes such as *CCL5*, *CCR5*, *BMP4*, *BMP7*, *CCDC88B* represent Chemokine signaling and bone morphogenetic protein (BMP) pathway activation, which contributes to vascular inflammation, immune activation, and arterial remodeling. Taken together, PCSF demonstrates a broad multi-pathway activation of inflammatory and immune regulatory mechanisms in HAECs following exposure to MPLs, potentially leading to endothelial dysfunction (HAECs) and progression of atherosclerosis, a main cardiovascular disease.

### 3.6. Comparative Transcriptomic Analysis Reveals Shared and Discordant Gene Regulation Between MPLs Treated Endothelial Cells and Human Atherosclerotic Plaques

To determine whether MPLs treated HAECs show gene expression patterns and features similar to human atherosclerosis, we compared the differentially expressed genes (DEGs) from MPLs treated HAECs with DEGs from a publicly available dataset (GSE120521) of human atherosclerotic plaques. The publicly available human atherosclerotic DEGs were derived by comparing stable plaques with unstable plaques. It is important to note that both stable and unstable plaques are atherosclerotic in nature, with unstable plaques representing the advanced stage of atherosclerosis. Therefore, comparing our MPLs-induced DEGs with DEGs from the public dataset may result in an underrepresentation of the full spectrum of gene expression changes induced by MPLs. As shown in **Fig. 10a**, out of 2,569 DEGs identified in atherosclerotic plaques and 1,637 DEGs from MPLs-treated HAECs, 457 DEGs were shared between MPLs treated HAECs and the human atherosclerotic plaques DEGs.

**Figure 10.**
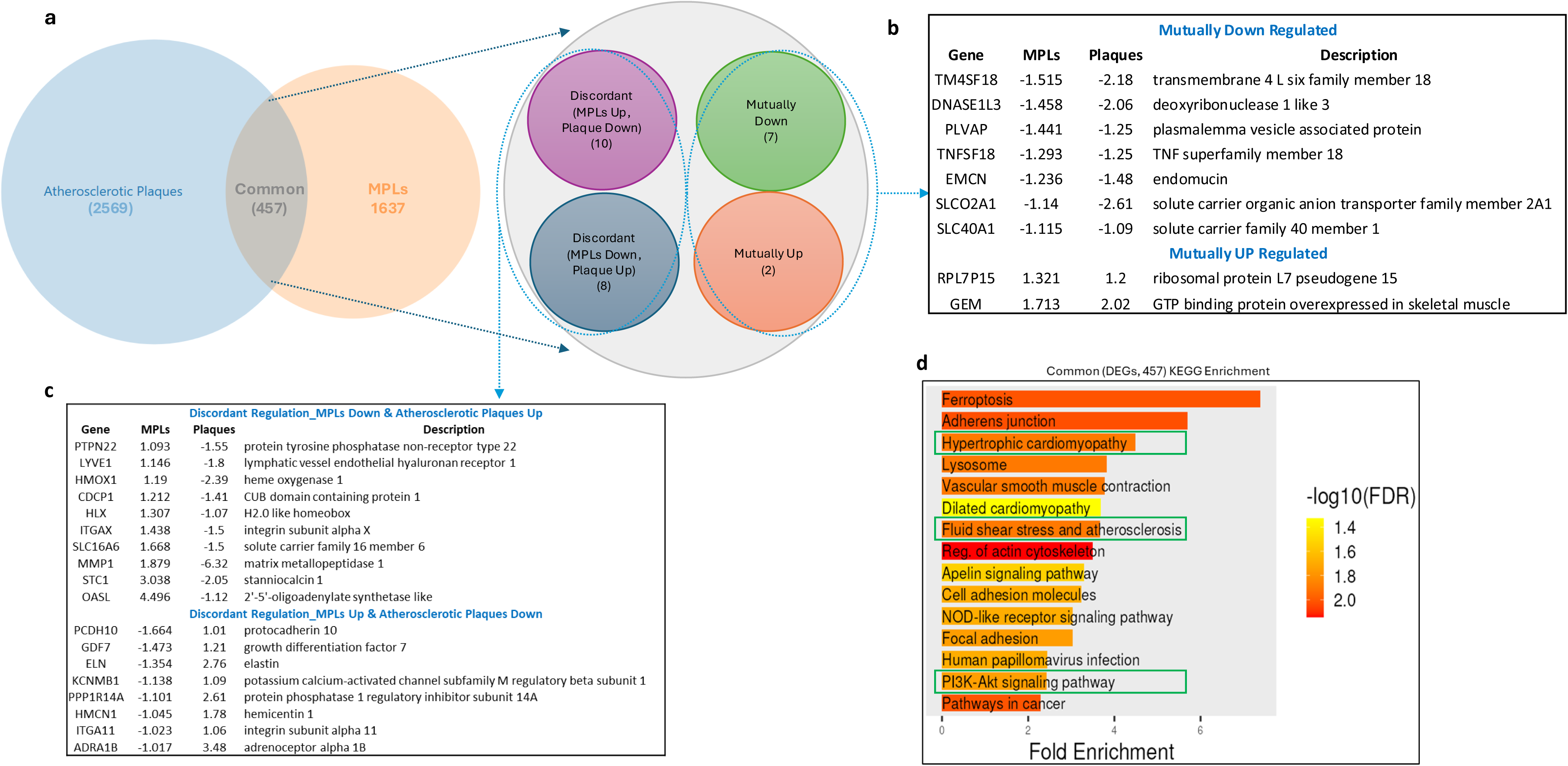
Comparative transcriptomic analysis of MPLs-treated HAECs and human atherosclerotic plaques reveals overlapping and distinct patterns of gene regulation. (a) Venn diagram showing the number of differentially expressed genes (DEGs) identified in MPLs-treated HAECs (n = 1,637) and the human atherosclerotic plaque dataset GSE120521 (n = 2,569), with 457 genes commonly altered in both conditions. (b) Classification of common DEGs into four regulatory categories: mutually downregulated (n = 7), mutually upregulated (n = 2), MPLs upregulated and plaque–downregulated (discordant, n = 10), and MPLs downregulated and plaque–upregulated (discordant, n = 8). (c) Tables listing genes from the discordant regulation categories with corresponding fold changes in MPLs treated and plaque datasets, and functional annotations. (d) KEGG pathway enrichment analysis of the 457 shared DEGs, highlighting enriched pathways including “Fluid shear stress and atherosclerosis,” “PI3K-Akt signaling,” “Focal adhesion,” “Regulation of actin cytoskeleton,” “Hypertrophic cardiomyopathy,” and “Lysosome.” Dot color represents −log₁₀(FDR), and dot size reflects fold enrichment.

Among the 457 shared DEGs, 7 genes (*TM4SF18, DNASE1L3, PLVAP, TNFSF18, EMCN, SLCO2A1,* and *SLC40A1*) were downregulated, and 2 genes (*RPL7P15* and *GEM*) were upregulated **(Fig. 10b)** in both datasets. In contrast, discordant gene regulation was observed in 18 genes **(Fig. 10c)**. Ten genes (*PTPN22, LYVE1, HMOX1, CDCP1, HLX, ITGAX, SLC16A6, MMP1, STC1, OASL*) were upregulated in the MPLs-treated group but downregulated in atherosclerotic plaques. 8 genes showed the opposite pattern, i.e., downregulated in the MPLs dataset but upregulated in the plaque dataset. These include *PCDH10, GDF7, ELN, KCNMB1, PPP1R14A, HMCN1, ITGA11,* and *ADRA1B*.

KEGG pathway enrichment analysis of the 457 common DEGs revealed overrepresentation of pathways relevant to vascular physiology and cell signaling **(Fig. 10d)**. Enriched pathways include Fluid shear stress and atherosclerosis, Hypertrophic cardiomyopathy, Dilated cardiomyopathy, PI3K-Akt signaling pathway, Regulation of actin cytoskeleton, Adherens junction, Lysosome, and Focal adhesion. Enrichment was quantified using fold enrichment and −log₁₀(FDR) values.

These data demonstrate a common gene expression pattern between MPLs-treated HAECs and human atherosclerotic plaques, highlighting both shared and divergent regulatory features.

### 3.7. MPLs Exposure induces epitranscriptomics changes leading to modification in canonical nucleotides in HAECs

RNA modifications, collectively known as the epitranscriptome, regulate many RNA-dependent cellular activities, including endothelial inflammation, gene expression, and, ultimately, the translation of mRNA into protein ^70–73^. MPLs had a clear effect on specific RNA modifications across the epitranscriptome. Across the 32 RNA modifications profiled, we classified each modification as significantly upregulated (fold change > 1.2, p < 0.05 vs. Ctrl), significantly downregulated (fold change < 0.8, p < 0.05 vs Ctrl), no change (0.8–1.2), or not detected (no measurable signal), and present the proportion of modifications in each category in the pie chart. As shown in the panel of bar graphs **(Fig. 11a)**, significant modifications were seen in various RNA Ribonucleosides. Among the adenosine modifications, levels of 1-methyladenosine (m^1^A) and N6-threonylcarbamoyladenosine (t^6^A) were significantly elevated following MPLs exposure, whereas N6,2’-O-dimethyladenosine (m^6^Am) showed a modest but statistically significant (p < 0.05) decrease. Uridine derivatives such as 3-(3-amino-3-carboxypropyl)uridine (acp^3^U) and Pseudouridine (Ψ) exhibited significant increases in normalized signal intensity. Among cytidine modifications, both 5-methylcytidine (m^5^C) and 3-methylcytidine (m^3^C) were significantly upregulated following treatment. For guanosine-related modifications, N2-methylguanosine (m^2^G), 7-methylguanosine (m^7^G), and N2, N2-dimethylguanosine (m^2,2^G) displayed marked increases in signal intensity in MPLs-treated HAECs as compared to the control group. These shifts in signal intensities reflect quantifiable changes in the abundance of the respective modified nucleosides under MPLs exposure.

**Fig. 11.**
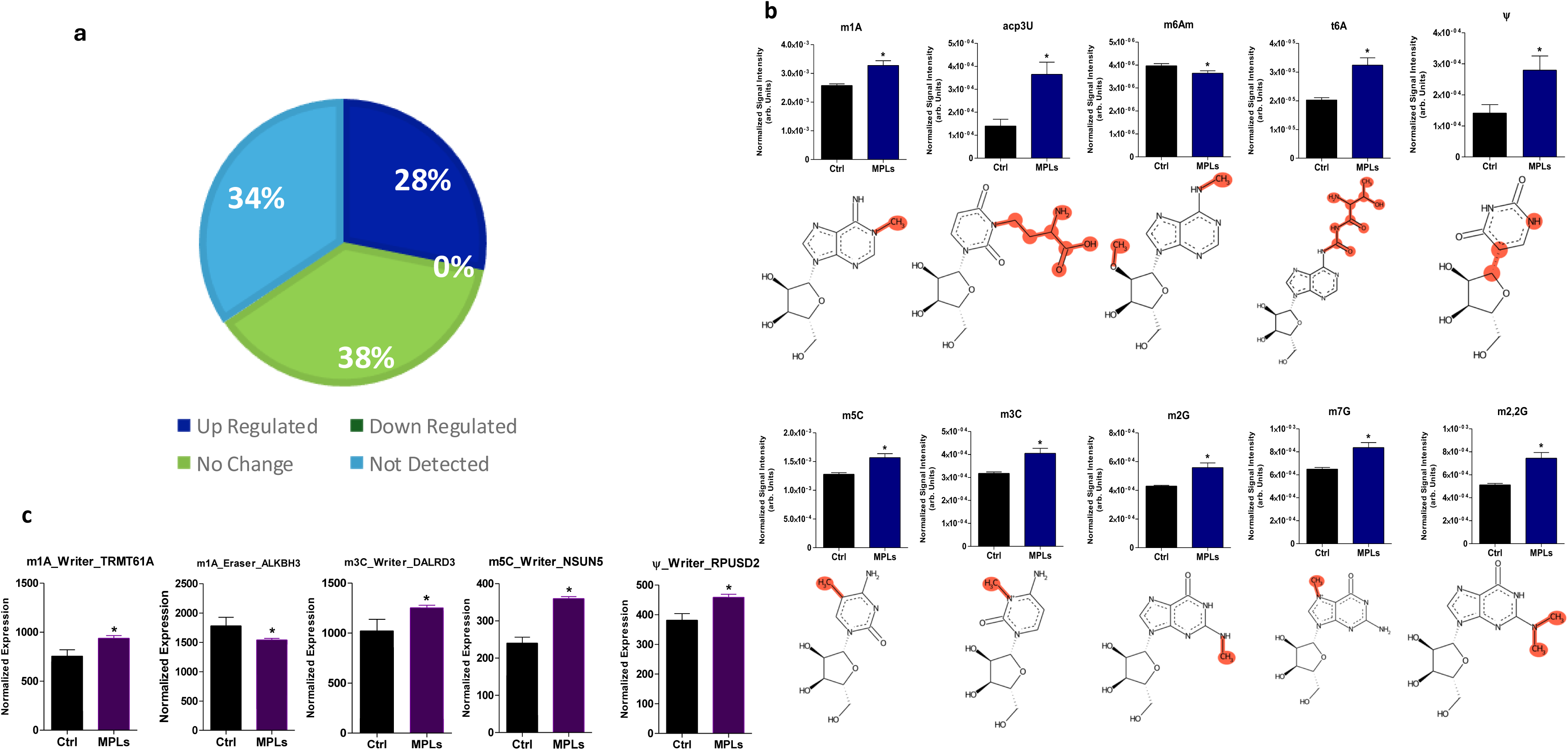
MPLs reshape RNA modifications and RNA modification machinery in HAECs. (a) RNA extracted from HAECs exposed to 80 nm MPLs (120 μg/mL) was quantified using UPLC-MS/MS. The pie chart summarizes the proportion of the 32 targeted RNA modifications that were significantly upregulated (fold change > 1.2, p < 0.05 vs. control), significantly downregulated (fold change < 0.8, p < 0.05 vs. control), no change (fold change 0.8–1.2), or not detected (no measurable signal). (b) Bar graph quantification of only significant (p < 0.05 vs Ctrl) measured from the total RNA extracted from MPLs exposed HAECs and the Ctrl group, with chemical structures and modified moiety highlighted in orange. Data are presented as mean ± SEM (n = 3). (C) Bar graphs represent key enzymes (genes) involved in RNA ribonucleoside modification in response to MPLs. Gene expression was quantified using RNA Sequencing, and normalized counts of three biological replicates were used per group. Data shown are mean ± SEM (n =3). * padj < 0.05 compared to the control group.

To determine how MPLs treatment affects epitranscriptome in HAECs, we quantified differentially expressed RNA modifications associated with writers, readers, and erasers from our transcriptomics data. We observed significant (p < 0.05) upregulation of m^1^A modification writers, *TRMT61A*, and *downregulation of the eraser, ALKBH3* (**Fig. 11c)**. Among m^5^C related enzymes, the writer *NSUN5* showed significant (p < 0.05) increased expression **(Fig. 11c)**. For m^3^C, the writer (cofactor) *DALRD3* was significantly (p < 0.05) upregulated **(Fig. 11c)**. For Ψ, our transcriptomics data indicated significant (p < 0.05) increased expression in the writer *RPUSD2* **(Fig. 11c)**. These results indicate that MPLs exposure is associated with altered steady-state levels of multiple RNA modifications and their regulatory enzymes. As these are steady-state measurements, they describe the epitranscriptomic state of MPLs-treated HAECs but do not, by themselves, establish downstream functional consequences.

### 3.8. LC-MS-Based Untargeted Metabolomics Reveals Distinct Metabolic Signatures in MPLs–Treated HAECs

A total of 3132 features were identified using untargeted LC-MS metabolomics following HAECs exposure to MPLs. PCA revealed clear group separation between MPLs and the Ctrl group, with 31.8% separation on PC1 and 16% on PC2, indicating a clear effect of MPLs on HAECs’ metabolites **(Fig. 12a)**. Heatmap of top 50 ranked feature demonstrated strong within group consistency and clear separation between MPLs and Ctrl group **(Fig. 12b)**. To identify differential metabolites (DMs) between MPLs and the Ctrl group, we used a t-test, which revealed a total of 268 DMs (FDR < 0.05) **(Fig. S1a)**. Differential metabolites obtained from the t-test output were annotated in SIRIUS. A list of differentially abundant features (FDR < 0.05) was identified between MPL-treated and control HAECs (Table S3). The putative annotated set include enrichment of oxidized and inflammatory phospholipid species that are consistent with proatherogenic lipid remodeling. Among them three truncated oxidized phosphatidylcholines (OXPCs) including 9-oxononanoyl PC (8256p), 8-oxooctanyl PC (7660p) and 5-oxo-pentanoyl PC (6905p) were significantly increased in MPLs group **(Fig. 12c-e)**. These species belong to the truncated aldehyde/oxo OxPC family structurally related to POVPC and PGPC, canonical components of oxidized LDL. Furthermore, lysophospholipid species were significantly increased in MPLs group, such as LysoPE(22:5) (5899p), PE(20:4/0:0) (5689p), 1-Hexadecanoyl-sn-glycero-3-phosphoethanolamine (6185p), 1-Oleoyl-sn-glycero-3-phosphoethanolamine (6491p), and 1-Stearoylglycerophosphoserine (LysoPS 18:0) (8005p). We also observed enrichment of arachidonoyl-containing PE (5689p), suggesting increased availability of eicosanoid precursors **(Fig. S1b-g)**. Acyl-acetyl phosphocholine species (6632p) which are structurally related to platelet activating factor (PAF) axis was also increased. Finally, we also observed increase in Phosphocholine (704p) **(Fig. S1h-i)**. Additionally, using a volcano plot (FDR < 0.05 and fold-change ≥ 2.0), we identified that 43 significantly decreased and 137 were increased in MPLs group relative to Ctrl **(Fig. S1j).**

**Figure 12.**
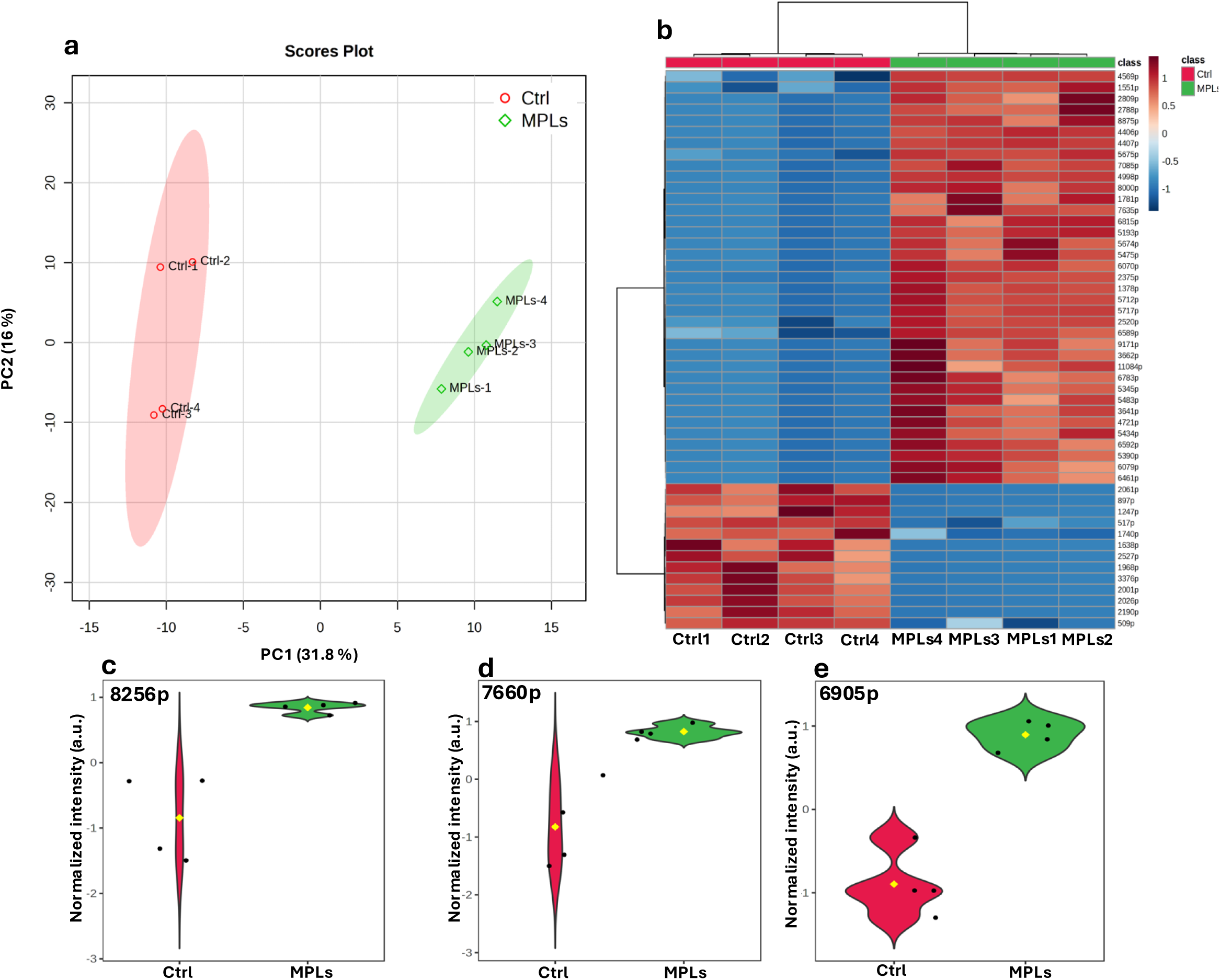
Untargeted LC-MS-based metabolomics reveals distinct metabolic profiles in MPLs–treated HAECs: (a) Principal Component Analysis (PCA) score plot showing clear separation between MPLs and Ctrl HAECs based on principal component 1 (PC1: 31.8%) and principal component 2 (PC2: 16%). (b) Heatmap of the top 50 differentially abundant metabolite features (FDR < 0.05) showing relative abundance across samples. Values are displayed as auto-scaled intensities (z-scores); red indicates higher and blue indicates lower relative abundance. (c-e) Representative violin plots of significantly increased oxidized phosphatidylcholine species in MPLs group, including 8256p (9-oxononanoyl PC), 7660p (8-oxooctanoyl PC), and 6905p (5-oxo-pentanoyl PC). Data are shown as normalized relative intensity (arbitrary units, a.u.). Yellow diamonds indicate group means; black dots represent individual biological replicates.

## 4. Discussion

Microplastics are ubiquitous environmental pollutants and are recognized as a threat to the ecosystem and human health, with growing evidence linking exposure to cardiovascular disease risk, such as atherosclerosis ^74, 75^. However, the uptake and routes of polystyrene microplastics (MPLs), their intracellular co-localization, and their mechanism of action are poorly understood in Human Aortic Endothelial Cells (HAECs). Therefore, we examined how MPLs are internalized by HAECs and the molecular pathways that may promote atherosclerosis by integrating transcriptomics, epitranscriptomics (RNA modifications), and metabolomics. HAECs line the interior of blood vessels, help maintain vascular homeostasis and functional regulation, and thus play a crucial role in the development of atherosclerosis. This study found that polystyrene microplastics (MPLs) can enter HAECs via multiple endocytic pathways, including clathrin-dependent endocytosis and macropinocytosis and through various channels. MPLs significantly induce inflammatory responses, particularly by activating the NF-κB signaling pathways. Subsequent luciferase reporter assays demonstrated that MPL treatment significantly increased NF-κB activity compared to the control, further confirming MPL’s role in inducing NF-κB transcriptional activation. Furthermore, MPLs treatment dysregulated multiple metabolite levels in HAECs, which are closely related to various disease processes. Simultaneously, MPLs exposure also led to abnormalities in various RNA modifications, suggesting changes in gene expression regulation. Further comparison of transcriptomic data from MPLs-treated HAECs and human atherosclerotic plaques revealed common dysregulation across multiple pathways, particularly those involved in vascular physiological regulation and cell signaling. Our studies reveal that HAECs can internalize MPLs, leading to multiple dysregulations in the transcriptome, epitranscriptome, and metabolic network, indicating that MPLs exposure may pose potential cardiovascular hazards to human health.

Our findings are the first to demonstrate that, after cellular uptake, MPLs are localized to mitochondrial, endolysosomal, and lysosomal compartments of human aortic endothelial cells. Consistent with this co-localization, our transcriptomics data showed disruption of mitochondrial genes, including decreased expression of mitochondrially encoded electron transport and oxidative phosphorylation genes (*MT-ND1/2/3/4/5/6, MT-CO1/2/3, MT-CYB, MT-ATP6*) and mitochondrial RNAs, suggesting impaired respiratory capacity and a shift towards MPLs induced redox stress. This is biologically meaningful, as MPLs are reported to disrupt mitochondrial homeostasis ^76–78^, drive endothelial cell senescence, and impair endothelial function via oxidative stress signaling, including reduced eNOS and altered redox pathways ^79^, thereby linking mitochondrial disruption to vascular pathology. MPLs have been reported to cause endothelial inflammation and adhesion molecule release, which are key steps in atherosclerosis, suggesting that MPLs targeting mitochondria may activate endothelial cells and recruit leukocytes ^80, 81^. Other studies have shown that polystyrene microplastics trigger ROS-mitochondrial pathways such as mitophagy, which may amplify inflammatory stress responses ^82, 83^. Together the apparent co-localization of MPLs with mitochondria and the mitochondrial gene changes in our transcriptomic data suggest the MPLs associate with and alter mitochondrial gene expression in HAECs, which may in turn contribute to endothelial dysfunction and hence atherosclerosis. Although endothelial cells generate most of their ATP through glycolysis and contain relatively few mitochondria, mitochondrial oxidative phosphorylation remains essential for endothelial homeostasis, and the electron transport chain, particularly Complex I, is a recognized determinant of endothelial redox balance and activation in atherogenesis ^84–87^. In this context, the coordinated suppression of Complex I and other ETC transcripts we observed is consistent with impaired endothelial mitochondrial function, positioning mitochondria as a plausible node linking MPLs exposure to endothelial dysfunction.

Additionally, MPLs co-localization was observed with endolysosomal and lysosomal compartments, consistent with previous work showing that polystyrene particles often accumulate in lysosomes and are processed via lysosomal-mediated exocytosis ^88, 89^. Previously, lysosome-centered handling of MPLs has been linked to broader remodeling of cellular programs, including lipid metabolism and immune activation in macrophage systems ^90^. MPLs’ interaction with lysosomes in our study is also important, as lysosomal lipid handling is central to vascular inflammation and plaque biology.

This study investigated the uptake pathways of MPLs in HAECs, to elucidate their internalization mechanisms and facilitate the development of corresponding intervention strategies. We found that HAECs internalize MPLs through macropinocytosis, clathrin-mediated endocytosis, and caveolae-mediated endocytosis, indicating that MPLs can enter cells via multiple endocytic pathways. Specifically, chlorpromazine significantly inhibited the uptake of MPLs by interfering with clathrin dissociation, thereby blocking the clathrin-mediated endocytic pathway ^91, 92^, suggesting its crucial role in MPLs internalization. Treatment with amiloride hydrochloride significantly reduced the cellular uptake of MPLs. This drug inhibits the ENaC pump and interferes with the Rac1/Cdc42 signaling pathway, thereby inhibiting macropinocytosis ^58, 93^, suggesting that macropinocytosis is also an important uptake pathway. Ebselen, a multi-target seleno-organic compound that inhibits mammalian H⁺/K⁺-ATPase ^63^ thereby disrupting endosomal/vesicular pH regulation ^64^ and also reacts with thiol-dependent enzymes including cysteine proteases[NEWand nocodazole, which hinders vesicle transport by inhibiting microtubule polymerization ^94, 95^, both significantly reduced MPLs uptake, further supporting the involvement of endocytic pathways.

Furthermore, the chloride channel blockers N-phenylanthranilic acid and anthracene-9-carboxylic acid interfere with endosomal acidification ^55, 66, 96, 97^, while amlodipine affects the dephosphorylation of endocytosis-related proteins by inhibiting calcium influx. Chloride channels play a role in neutralizing endosomal acidification ^98^. Endosomes naturally have a pH of 6.5–4.5, and anything outside of that range has the potential to disrupt the sorting capabilities of endosomes and, therefore, endocytic processes. Although ion balance and endosomal pH regulation appear to contribute to MPLs internalization, direct inhibition of the vacuolar H⁺-ATPase with bafilomycin A1 did not significantly reduce uptake, indicating that endosomal acidification may not essential for the initial internalization step but likely acts later in endosomal maturation and cargo sorting. Collectively, this study systematically demonstrates that MPLs can enter HAECs via multiple endocytic pathways, and that these pathways depend on cellular processes such as clathrin dynamics, cytoskeletal transport, ion channel function, and endosomal acidification, providing experimental evidence for a deeper understanding of the mechanisms of microplastic-induced endothelial toxicity.

Our transcriptome analysis showed that endocytosis and vesicular transport pathways were significantly enriched in HAECs treated with MPLs. In this study, the Rab GTPases, actin-binding proteins, and SNX genes that were significantly regulated in MPLs-treated HAECs play major roles in cellular uptake. Rab GTPases localize to various endosomal structures and regulate endosomal function by recruiting various factors ^99^. Disruption of genes encoding actin and many actin-binding proteins has been shown to block the uptake of endocytic markers ^100^. SNXs also play pivotal roles throughout the endocytic trafficking pathway, including endocytosis, endosomal sorting, and endosomal signaling. Thus, the upregulation of Rab GTPases, actin-binding proteins, and SNX genes in MPLs-treated HAECs, as revealed by RNA-seq analysis in this study, adds another layer of evidence that MPLs are internalized by HAECs.

Our transcriptome analysis revealed differential gene expression patterns in HAECs exposed to MPLs. Inflammation plays a decisive role in the development and progression of atherosclerosis^101–104^. Damaged endothelial cells secrete pro-inflammatory cytokines, chemokines, and cell adhesion molecules, thereby promoting the development of atherosclerosis^101–104^. We hypothesized that MPLs uptake can stimulate inflammatory pathways in ECs, as MPLs act as foreign bodies in living systems, prompting an immune response. Our findings show that MPLs activated NF-κB signaling in HAECs and the subsequent demonstration that NF-κB had higher luciferase activity in MPLs-treated HAECs compared with control cells are consistent with previous studies. High MPLs concentrations induced inflammation in the small intestine by activating the TLR4 signaling pathway ^105^. Neural stem cells and neurons exposed to various concentrations of MPLs showed upregulated pathways involved in neuroinflammation, innate and adaptive immunity, cell migration, proliferation, extracellular matrix remodeling, and cytoskeleton structures ^106^. It has also been reported that MPLs induced the onset of the pyroptosis signaling pathway in human vascular endothelial cells ^107^. In this study, the upregulation of genes associated with biological processes related to host defense and external stimulus response upon exposure of HAEC cell lines to MPLs, as revealed by gene ontology analysis, supports our hypothesis that MPLs can indeed evoke an inflammatory response.

MPLs induce inflammatory responses by TNF-α signaling stimulation ^108^. Mice endothelial cells exposed to MPLs evoke inflammatory pathways, including the TNF-α NF-κB signaling pathway, to express inflammatory proteins ^81^. Increased levels of inflammatory markers, including NF-κB, IL-6, TNF-α, and CD68, were observed when MPLs accumulated in adipose tissue ^109^. Consistent with these previous studies, MPLs-treated HAECs in this study revealed TNF-α signaling via NF-κB as an enriched pathway, along with many other pathways. As stated previously, the transcriptional activation of TNF-α via the NF-κB signaling pathway in this study was also confirmed by a significant increase in relative luminescent units in MPLs-treated HAEC cell lines, adding another compelling piece of evidence to the role that MPLs used in this study influence inflammation and their potential role in disrupting endothelial cells. MPLs trigger pronounced pathological changes in epithelial-mesenchymal transition in mice ^110^, including increased migration rate, decreased cell adhesion, and cytoskeletal rearrangement. Fibronectin, a reliable marker of epithelial-mesenchymal transition, was significantly reduced in MPLs-treated human gingival fibroblast (HGF) cells compared to untreated control cells ^111^. Thus, our finding that MPLs-treated HAECs show downregulated gene sets, such as epithelial-mesenchymal transition, is consistent with previous studies and points to the role of MPLs in compromising cellular function in HAECs. Our observation of a downregulated gene set of E2F transcription factor targets is consistent with a previous study, which found that E2F targets were downregulated in MPLs-treated HGF cells ^111^. E2Fs are important regulators of genes required for cell cycle progression and play an integral role in controlling cell proliferation ^112^. In this study, the downregulation of genes involved in cell structure and morphogenesis, cell adhesion, and cell migration is expected upon exposure of HAEC cell lines to MPLs. Adverse effects of MPLs on living cells have been reported in the literature, including reduced cell proliferation, significant changes in cell morphology, and decreased metabolic activity ^113, 114^.

A clear pro-atherogenic state induced by MPLs in HAECs is evident from our curated figure showing upregulated pro-atherosclerotic and downregulated anti-atherosclerotic genes; i.e., MPLs push HAECs into an active endothelium state, a state known to seed early atherosclerosis, and simultaneously downregulate genes for protective and lipid-handling systems ^115^, thereby promoting atherosclerosis. This aligns with previous studies showing that polystyrene MPLs can trigger endothelial activation and inflammation ^81, 116^. The upregulation of *CCL5* and chemokine-related genes is very important because chemokines drive leukocyte recruitment and lesion formation in atherosclerosis ^117, 118^. Type 1 interferon signaling, which can promote inflammation and adhesion signaling ^119, 120^, was also upregulated in our study. Together, our curated genes suggest that MPLs may accelerate the risk of atherosclerosis by turning on immune recruitment, interferon, and inflammatory programs, while tuning down anti-atherosclerotic programs, thereby creating a vascular environment that is more inflammation-prone and less resilient ^121^.

We further used comparative transcriptomics analysis to reveal the commonalities and differences in gene regulation between MPLs-treated endothelial cells and human atherosclerotic plaques. By integrating our MPLs-treated HAECs transcriptomic data with publicly available transcriptomic data on atherosclerosis plaques, we have established that MPLs treated HAECs closely parallel atherosclerosis plaque at the whole-genome expression level in terms of gene enrichment in Fluid shear stress and atherosclerosis pathway, Hypertrophic cardiomyopathy pathway, Dilated cardiomyopathy pathway, PI3K-Akt signaling pathway, Regulation of actin cytoskeleton, Adherens junction, Lysosome, and Focal adhesion. All these pathways play crucial roles in maintaining endothelial integrity, and dysregulation of their genes can lead to endothelial dysfunction, inflammation, and atherosclerotic disease progression. It is important to note that the expression profile derived from cell lines was compared with that of atherosclerotic plaques in publicly available data, therefore, the observed divergence in gene expression direction between the two datasets is to be expected.

Post-transcriptional RNA modifications have recently emerged as critical regulators of gene expression, and they influence RNA processing, metabolism, and other critical pathways ^122, 123^. We employed mass spectrometry (MS) to identify and quantify RNA modifications in HAECs after exposure to MPLs. Consistent with a recent study, which detected and quantified twenty dysregulated modifications from small RNA and nine modifications from mRNA when various mouse tissues were exposed to MPLs, our results revealed significant modifications of RNA ribonucleosides within HAECs after exposure to MPLs. The statistically significant changes most notably observed in N^1^-methyladenosine (m^1^A), N^6^,2ʹ-O-dimethyladenosine (m^6^Am), N^2^-methylguanosine (m^2^G), N^7^-methylguanosine (m^7^G), N^2^,N^2^-dimethylguanosine (m^2,2^G), N^3^-methylcytidine (m^3^C), N^5^-methylcytidine (m^5^C), pseudouridine (Ψ), 3-(3-amino-3-carboxypropyl)uridine (acp^3^U), and N^6^-threonylcarbamoyladenosine (t^6^A) indicated the dysregulation of these modifications. Previous studies have implicated several of the dysregulated RNA modifications found in our study to have a role in the etiology and progression of diseases, including cardiovascular diseases. Adenosine ribonucleoside modification, i.e., m6A, promotes endothelial inflammatory responses and facilitates the formation of AS plaque, and its inhibition reduces endothelial inflammation and mitigates AS progression in mice ^124^. Dysregulation of t6A modification and its biosynthetic enzymes (*OSGEP*/*OSGEPL1*) has been associated with the development and progression of several malignancies, including hepatocellular carcinoma ^125^. Elevated plasma Ψ concentrations have been observed in individuals with heart failure compared to healthy controls ^126^. Pseudouridylation can be induced under stress conditions ^127^. Levels of acp^3^U are high in supernatants from breast carcinoma cells ^128^. Decreased level of m^7^G in the mRNA of ischemic tissues, implies an association between m^7^G and vascular disease ^129^. RNA methyltransferase *NSUN2*, which catalyzes m^5^C formation ^130^, has been shown to increase leukocyte adhesion to the endothelium ^131^, which is a hallmark of the inflammatory process. These previous studies provide hints of the hazardous effects of MPLs in HAECs, possibly endothelial dysfunction, and the progression of atherosclerosis.

Integrated transcriptomics–epitranscriptomics analysis reveals that MPLs exposure promotes a shift toward higher m^1^A RNA methylation in HAECs, characterized by TRMT61A upregulation (writer) and *ALKBH3* downregulation (eraser), together with a global increase in m^1^A. TRMT6/TRMT61A is a canonical methyltransferase complex that installs tRNA m^1^A58, a modification which is known to support tRNA structure and translational control ^132^. Evidence suggests that TRMT6/TRMT61A also installs m^1^A-specific sites on mRNA, with substrate selection guided by the GUUCRA motif and a local tRNA T-loop-like structure; hence, mRNA recognition by the *TRMT6*/*TRMT61A* complex relies on principles similar to those of canonical tRNA substrate recognition ^133^. In contrast, *ALKBH3* functions as tRNA m^1^A demethylase, and reduced *ALKBH3* activity is consistent with increased tRNA m^1^A level ^134^. *ALKBH3* also serves as a demethylase for m^1^A in mRNA ^135^. A study reported that 774 m^1^A peaks in *ALKBH3* knockout HEK293T cells ^135–137^. Therefore, the *TRMT61A*-up and *ALKBH3*-down configuration observed in our transcriptomics data, mechanistically supports m^1^A accumulation and, hence, suggests that MPLs shape endothelial cell fate by coordinated transcriptomic and epitranscriptomic reprogramming.

Notably, with the increase in m^1^A modification, we observed transcriptional upregulation of genes that are involved in endothelial inflammation and immune recruitment such as upregulation of chemokine and chemokine receptor genes (*CCL5*, *CXCL16*, *CXCR4*), immune interface and adhesion related genes (*ICAM2*, *CD44*, *CD276*), and a prominent interferon-stimulated gene module (*IFI27*, *IFI35*, *IFI44*, *IFI44L*, *IFI6*, *IFIH1*, *IFIT1*/*2*/*3*). This co-occurrence is consistent with endothelial cells adopting a pro-inflammatory state to promote leukocyte–endothelial interaction and amplify vascular immune signaling, a clear pattern of early events in atherogenesis. However, whether m1A directly regulates these specific transcripts cannot be determined from bulk measurements and would require transcript-resolved m1A mapping (e.g., m1A-seq). Previously, MPLs have induced endothelial activation with increased adhesion molecule expression and subsequent leukocyte adhesion ^81^. As RNA modifications such as m^6^A are broadly recognized as contributors to vascular pathology, including atherosclerosis, recent literature suggests that other RNA modifications are implicated in cardiovascular diseases ^73, 138^. Our findings (*TRMT61A*↑/*ALKBH3*↓; m¹A↑, *CCL5*↑, *CXCR4*↑, *ICAM2*↑, *IFI27*↑) therefore suggest a possible association between MPLs-induced epitranscriptomic changes and the pro-atherogenic transcriptional state of HAECs, which remains to be tested directly.

Similarly, we observed increased expression of DALRD3, a cofactor required for human METTL2-dependent installation of m^3^C in human tRNA ^139, 140^, along with increased expression of *RPUSD*2, a member of the *RPUSD2* family of pseudouridine synthases; family members, notably *RPUSD4*, catalyze Ψ formation in mitochondrial RNA, including 16S mt-rRNA and mt-tRNAs ^141^. Furthermore, the expression of *NSUN5*, an enzyme involved in installing m^5^C at C3782 of human 28S rRNA and influencing ribosome function, was also increased ^142^. Finally, we observed upregulation of various methyltransferase-associated genes (*METTL23*, *METTL27*) and TRMT112-related genes (*TRMT112P6*), consistent with a generalized increase in methylation and translation. Additional targeted work will be needed to define whether the observed increase in various other RNA modifications, such as m^7^G, is directly linked to the MPL-induced increase in the writers of these modifications.

Our LC-MS-based untargeted metabolomics study revealed unique metabolic characteristics in HAECs treated with MPLs, demonstrating coordinated biochemical remodeling of HAECs. These metabolomics changes provide a mechanistic basis to evaluate how MPLs may promote endothelial dysfunction and atherosclerosis. Metabolomics analysis revealed differences in metabolites between control cells and MPLs-treated HAECs, indicating that MPLs regulate metabolic pathways in HAECs. This shift in metabolic profile after HAECs were treated with MPLs is consistent with previous studies ^143–145^. Metabolic pathways are key regulators of endothelial cells’ functions, including angiogenesis and inflammation ^146^.

The complete list of differentially abundant metabolites is provided in Table S3. Among the identified metabolites, we observed an increase in truncated oxidized phosphatidylcholines that are established mediators of endothelial dysfunction in atherosclerosis. POVPC and PGPC-like OxPCs accumulate in oxidized LDL and atherosclerotic lesions and directly promote processes involved in atherosclerosis ^147, 148^. The increase in multiple truncated OxPC species such as 8256p, 7660p, 6905p in MPLs group suggests oxidative fragmentation of membrane phosphatidylcholine which is a central biochemical feature of early atherogenesis. Our study suggests activation of phospholipase A_2_-dependent membrane remodeling as we observed an increase in lysophosphatidylethanolamine (5899p, 5689p, 6185p, 6491p). Lipoprotein associated PLA_2_ accumulates within atherosclerotic plaque and serves as predictor of cardiovascular risk, hence linking lysophospholipids to vascular inflammation ^149, 150^. Importantly, higher levels of arachidonoyl-PE (5689p) suggest greater availability of arachidonic acid, which supports cyclooxygenase and lipoxygenase pathways that form pro-inflammatory eicosanoids linked to endothelial activation and the plaque progression ^151^. An increase in LysoPS 18:0 (8005p) suggests the presence of lipid environment that promotes inflammation. LysoPS is known to stimulate macrophage foam cell development and trigger inflammatory response associated with atherosclerosis ^152^, suggesting that MPLs induced metabolites changes in HAEC composition could promote immune activation. Additionally, the presence of several glycerophospholipid candidates is important, as oxidized and remodeled phospholipids play a role in endothelial activation and atherosclerosis ^153, 154^.

Acyl-acetyl phosphocholine is an acyl analogue of platelet-activating factors that can bind to and activate the platelet-activating factor receptor (PAFR) ^155–158^. Our results show that the levels of acyl-acetyl-phosphocholine (6632p) were significantly elevated following MPL treatment, suggesting activation of the PAF signaling pathway ^155–158^, which is present in human atherosclerotic plaques and plays a role in vascular inflammation ^159^. Simultaneously, increased phosphocholine suggests accelerated phosphatidylcholine metabolism and oxidative breakdown of lipids. The phosphocholine headgroup in oxidized phospholipids is targeted by innate immune components such as C-reactive protein and natural antibodies, thereby connecting the buildup of oxidized phospholipids to inflammatory responses in blood vessels ^160^. PAFR activation has been shown to induce NF-κB signaling through G protein–coupled pathways, promoting IκBα degradation and nuclear translocation of NF-κB subunits ^161–164^, thereby activating the expression of NF-κB-dependent inflammatory and angiogenic mediators. Oxidized phospholipids and certain lysophospholipids are established regulators of inflammatory signaling in endothelial cells. In endothelial cells, lysophosphatidylcholine and lysophosphatidic acid can activate NF-κB and induce the expression of inflammatory chemokines and adhesion molecules ^165, 166^. Therefore, the significant increase in oxidized phospholipids and lysophospholipids induced by MPL exposure can activate the NF-κB signaling pathway, thereby establishing a link between MPL-driven lipid alterations and the expression of inflammatory genes in human aortic endothelial cells. Together, this evidence suggests that exposure to MPLs promotes the buildup of oxidized phospholipids, activation of phospholipases and an increase in eicosanoid precursors in HAECs, mirroring fundamental biochemical events seen at the onset of atherosclerosis.

Several limitations should be considered when interpreting our results. A limitation of this study is that the physicochemical properties of the polystyrene microplastics were not characterized within the complete cell culture medium used for the exposure experiments. Although SEM and FTIR confirmed the surface morphology, particle size, polymer type, and chemical composition of the microplastics, these analyses did not assess changes such as hydrodynamic particle size, aggregation state, surface charge, or protein corona formation induced by the culture medium. Interactions between the microplastics and components of the medium (e.g. salts, serum proteins) could alter particle behavior, sedimentation characteristics, and the actual dose delivered to the cells. Consequently, the physicochemical state of microplastics during cell exposure may differ from that observed under characterization conditions involving aqueous suspensions or dry samples. Future research should evaluate parameters such as particle size distribution, polydispersity, zeta potential, aggregation kinetics, and protein corona formation within the complete exposure medium under relevant concentration and incubation conditions. For our epitranscriptomic analysis, we quantified steady-state RNA modifications and enzyme expression but did not functionally test their downstream effects, which will require transcript-resolved mapping (m1A-seq/m5C-seq), enzyme knockdown or overexpression, and an assessment of the impacts on the transcriptome and metabolome. In addition, our metabolomic analysis is also constrained by annotation coverage, as only a subset of significant features could be putatively identified. Furthermore, mitochondrial respiration, membrane potential, and ROS were not directly measured, so the proposed link between MPLs and impaired endothelial mitochondrial function remains inferential and requires functional validation in future work.

## 5. Conclusion

In summary, this study is the first to show that human aortic endothelial cells can internalize polystyrene MPLs via multiple endocytic pathways, an uptake that is accompanied by inflammatory activation, metabolic disturbances, abnormal RNA modifications, and transcriptomic dysregulation. Our results suggest that microplastic exposure may be a potential risk factor for atherosclerosis by disrupting vascular endothelial homeostasis, providing a mechanistic basis for understanding how MPLs impair endothelial integrity and lead to adverse health consequences. By linking MPLs uptake to integrated transcriptomic, epitranscriptomic, and metabolomic reprogramming in human aortic endothelial cells, this study offers an early, hypothesis-generating framework and candidate molecular signatures (e.g., inflammatory and RNA-modification pathways) that could help future biomarker discovery and inform risk assessment for microplastic-associated cardiovascular disease. Realizing this translational potential will require direct functional endothelial measurements, alongside validation in donor-diverse primary cells and in vivo models, and exposure studies at environmentally relevant concentrations.

## Supporting information

https://uncg-my.sharepoint.com/:b:/g/personal/a_khan10_uncg_edu/IQD8uApsNxSlR6iC2uTqTmRQATMu_BkkH7tDobksmyKinBQ?e=JIFBOp

## Conflict of interest

None declared.

## Acknowledgments

This work was supported by a research grant from UNC Greensboro (127313).

## CRediT authorship contribution statement

**Ajmal Khan**: Conceptualization, Methodology, Investigation, Formal analysis, Data curation, Visualization, Writing – original draft, Writing – review & editing. **Grant Koher**: Investigation. **Tariq Khan**: Investigation. **Kyla Grant**: Investigation. **Sarah Hudson**: Investigation. **Gordon Zheng**: Formal analysis. **Ho Young Lee**: Investigation, Formal analysis. **Warren S. Vidar**: Investigation, Formal analysis. **Frank Morales-Shnaider**: Investigation. **Jinlan Chen**: Investigation. **Kwame A. Darfour-Oduro**: Writing – review & editing. Ramji Bhandari: Supervision. **Xuewei Zhu**: Resources, Supervision. **Kerui Wu**: Supervision. **Norman Chiu**: Investigation, Resources, Supervision. **Zhenquan Jia**: Conceptualization, Supervision, Project administration, Funding acquisition, Writing – review & editing.

## Figure legends

**Supplementary Fig. S1. Differential metabolite features in MPLs-treated HAECs.**

(a) Univariate statistical distribution of all detected LC–MS features comparing MPLs-treated and control (Ctrl) HAECs. Purple points indicate significant features (FDR < 0.05); gray points indicate non-significant features. (b–i) Representative violin plots of selected significantly increased metabolites in MPLs-treated HAECs, including LysoPE(22:5) (5899p), 1-Hexadecanoyl-sn-glycero-3-phosphoethanolamine (6185p), 1-Oleoyl-sn-glycero-3-phosphoethanolamine (6491p), LysoPS 18:0 (8005p), PE(20:4/0:0) (5689p), 1-Pentadecanoyl-2-acetyl-sn-glycero-3-phosphocholine (6632p), and phosphocholine (704p). Data are shown as normalized relative intensities (arbitrary units, a.u.). Yellow diamonds denote group means; dots represent biological replicates. (j) Volcano plot of MPLs-treated versus Ctrl HAECs showing log₂(fold change) versus –log₁₀(p-value). Red and blue points indicate significantly increased and decreased features, respectively (FDR < 0.05; |FC| ≥ 2); gray points indicate non-significant features.

**Table S1.** Uptake assay inhibitors. List of channels blockers and internalization pathways inhibitors. Full name, abbreviation, concentration, and the role of potential inhibitors in MPLs uptake.

**Table S2.** Parameters used of Mzmine peak picking analysis of MS raw data.

**Table S3.** Differential metabolite features identified in MPLs-treated HAECs by untargeted LC–MS metabolomics (FDR < 0.05).

## Notes

### Competing Interest Statement

The authors have declared no competing interest.

### Summary of Updates

The manuscript has been updated as follows: 1. Added cell viability data. 2. Corrected grammar and improved sentence structure throughout the manuscript. 3. Expanded and clarified the Discussion section. 4. Added a Limitations section. 5. Added an Author Contributions (CRediT) statement.

