## Supplementary material for "Polystyrene microplastic uptake drives Inflammatory, Epitranscriptomic, and Metabolic Reprogramming in Human Aortic Endothelial cells": https://uncg-my.sharepoint.com/:b:/g/personal/a_khan10_uncg_edu/IQD8uApsNxSlR6iC2uTqTmRQATMu_BkkH7tDobksmyKinBQ?e=JIFBOp

**Supplementary Fig. S1. Differential metabolite features in MPLs-treated HAECs.** (a) Univariate statistical distribution of all detected LC–MS features comparing MPLs-treated and control (Ctrl) HAECs. Purple points indicate significant features (FDR < 0.05); gray points indicate non-significant features. (b–i) Representative violin plots of selected significantly increased metabolites in MPLs-treated HAECs, including LysoPE(22:5) (5899p), 1-Hexadecanoyl-sn-glycero-3-phosphoethanolamine (6185p), 1-Oleoyl-sn-glycero-3-phosphoethanolamine (6491p), LysoPS 18:0 (8005p), PE(20:4/0:0) (5689p), 1-Pentadecanoyl-2-acetyl-sn-glycero-3-phosphocholine (6632p), and phosphocholine (704p). Data are shown as normalized relative intensities (arbitrary units, a.u.). Yellow diamonds denote group means; dots represent biological replicates. (j) Volcano plot of MPLs-treated versus Ctrl HAECs showing $\log_2$(fold change) versus $-\log_{10}$(p-value). Red and blue points indicate significantly increased and decreased features, respectively (FDR < 0.05; |FC| ≥ 2); gray points indicate non-significant features.

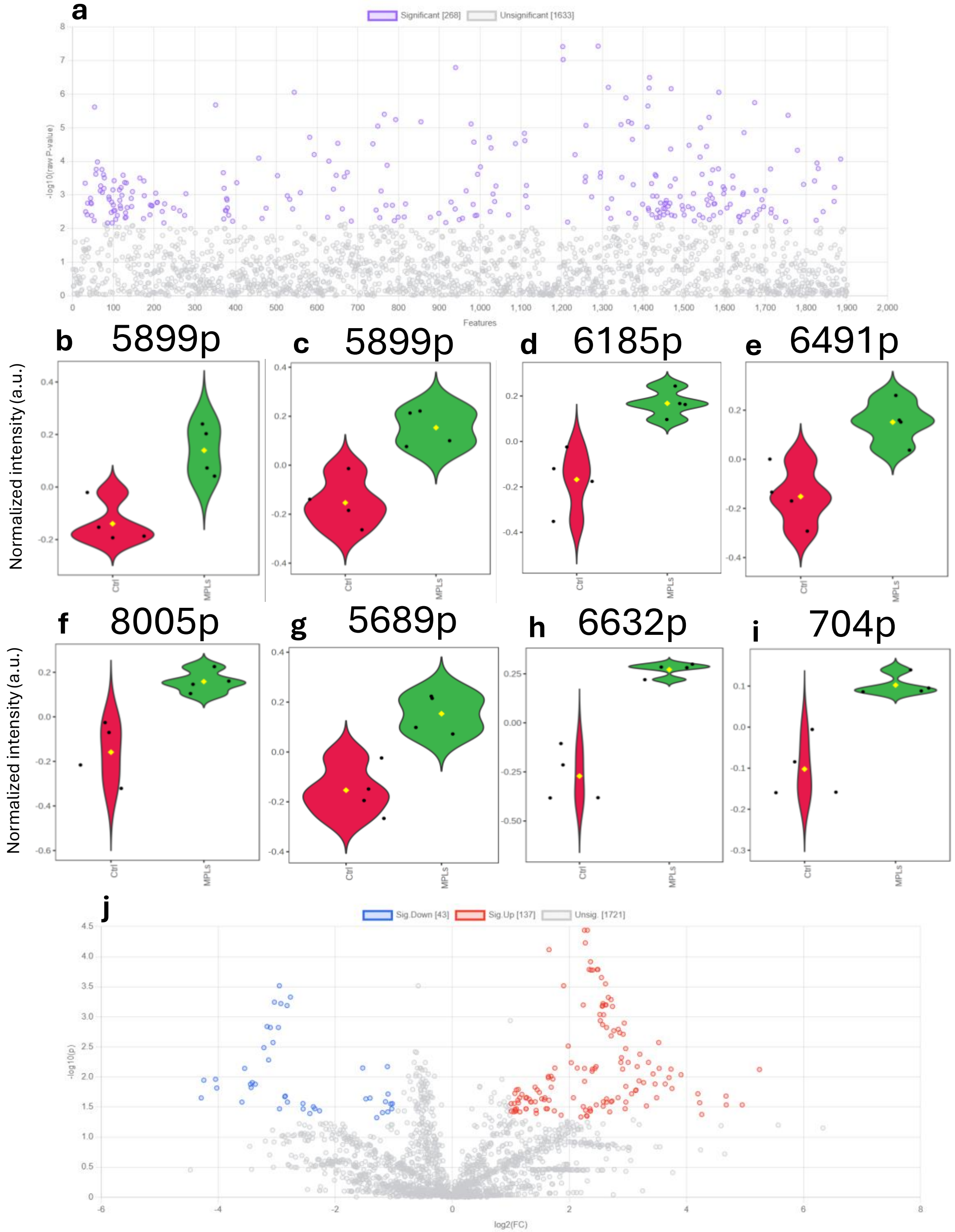

**Table S1.** Uptake assay inhibitors. List of channels blockers and internalization pathways inhibitors. Full name, abbreviation, concentration, and the role of potential inhibitors in MPLs uptake.

| Inhibitor | Abbrev | Conc | Role |
|---|---|---|---|
| **Bafilomycin** | Bf | | Bafilomycin A1 reversibly inhibits late-phase autophagy. By inhibiting H+-ATPase, bafilomycin A1 prevents re-acidification of synaptic vesicles following exocytosis and thereby blocks fusion of autophagosomes and lysosomes. Bafilomycin A1 can also inhibit cell proliferation [48-54] |
| **Chlorpromazine HCL** | Chl | 10 μM | Suppresses clathrin disassembly [55, 56] |
| **Amiloride Hydrochloride Dihydrous** | Amil | 50 μM | Inhibits micropinocytosis: blocks Na+/H+ exchanger pump [56-58] |
| **Anthracene-9-Carboxilic Acid** | Ant | 100 μM | Ion channel blocker (Cl−) [59] |
| **Amiodarone Hydrochloride** | Amio | 10 μM | Non-selective ion channel blocker [60] |
| **Genstein** | Gen | 200 μM | Inhibits tyrosine kinase receptors [55] |
| **Niflumic Acid** | Nif | 10 μM | |
| **Cobalt (II) Chloride** | Co | 2 mM | Ion channel blocker ($Ca^{2+}$), non-selective/inorganic [61] |
| **Amlodipine** | Aml | 10 μM | Ion channel blocker ($Ca^{2+}$), L-type/dihydropyridine [62] |
| **Ebselen** | Eb | 15 μM | Inhibits mammalian H+, K+-ATPase [63, 64] |
| **4-Aminopyridine ~98%** | 4-AP | 5 mM | Ion channel blocker (K+) [65] |
| **N_Phenlanthranilic Acid** | N-Ph | 0.1 mM | Ion channel blocker (Cl−) [66] |
| **Barium Chloride Anhydrous** | Ba | 350 μM | Ion channel blocker (K+) [65] |
| **Cesium Chloride 99%** | Cs | 1mM | Ion channel blocker (K+) [67] |
| **Phenylglyoxal** | Phen | 100 μg | Selective inhibitor of phagocytosis [68] |
| **Nocodazole** | Noc | 20 μM | Actin and microtubule disruptor [55] |
| **Mercury Chloride** | Hg | 50 μM | hAQPI Aquaporins [69] |
| **Cytochalasin A** | Cyt | 5 μg/mL | Actin disruptor [55] |
| **Copper Sulfate** | Cu | 100 μM | hAQP3 Aquaporins [69] |

**Table S2.** Parameters used of Mzmine peak picking analysis of MS raw data

| Module | Parameters Used |
|---|---|
| **A. Mass Detection (first, for MS1 data)** | |
| 1. MS Level Filter<br>2. Polarity<br><br>3. Spectrum Type<br>4. Mass Detector<br>5. Noise Level | MS1, level 1<br>positive and negative<br>(processed in separate batches)<br>centroid<br>centroid<br>5.0E3 |
| **B. Mass Detection (second, for MS2 data)** | |
| 1. MS Level Filter<br>2. Polarity<br><br>3. Spectrum Type<br>4. Mass Detector<br>5. Noise Level | MS2, level 2<br>positive and negative<br>(processed in separate batches)<br>centroid<br>centroid<br>100 |
| **C. Chromatogram Builder** | |
| 1. MS Level Filter<br>2. Polarity<br><br>3. Spectrum Type<br>4. Min. Consecutive Scans<br>5. Min. Intensity for Consecutive Scans<br>6. Min. Absolute Height<br>7. *m/z* Tolerance (scan-to-scan) | MS1, level 1<br>positive and negative<br>(processed in separate batches)<br>centroid<br>5<br>5.0E3<br>1.0E5<br>0.003 *m/z* or 5.0 ppm |
| **D. Local Minimum Feature Resolver** | |
| 1. MS1 to MS2 Precursor Tolerance (*m/z*)<br>2. Retention Time Filter<br>3. Minimum Relative Feature Height<br>4. Dimension<br>5. Chromatographic Threshold<br>6. Minimum Search Range RT<br>7. Minimum Absolute Height<br>8. Min. Ratio of Peak Top/Edge<br>9. Peak Duration Range<br>10. Minimum Scans (data points) | 0.003 m/z or 5.0 ppm<br>Use Tolerance, 0.20 min.<br>25%<br>Retention Time<br>70%<br>0.05<br>1.0E5<br>1.50<br>0.00 to 5.00<br>5 |
| **E. $^{13}C$ Isotope Filter** | |
| 1. *m/z* Tolerance (intra-sample)<br>2. Retention Time Tolerance<br>3. Monotonic Shape<br>4. Maximum Charge<br>5. Representative Isotope<br>6. Never Remove Feature with MS2 | 0.0015 *m/z* or 0.0 ppm<br>0.05 min<br>Checked<br>2<br>Most Intense<br>Checked |
| **F. Isotopic Peaks Finder** | |
| 1. Chemical Elements<br>2. *m/z* Tolerance (feature-to-scan)<br>3. Maximum Charge of Isotope *m/z*<br>4. Search in Scans | H, C, N, O, S, Cl, Br, P, F<br>0.002 *m/z* or 0.0 ppm<br>2<br>Single Most Intense |
| **G. Join Aligner** | |
| 1. *m/z* Tolerance (sample-to-sample)<br>2. Weight for *m/z*<br>3. Retention Time Tolerance<br>4. Weight for RT<br>5. Mobility Weight | 0.003 *m/z* or 5.0 ppm<br>2<br>0.10 minutes<br>1<br>1.0 |
| **H. Peak Finder (Gap Filling)** | |
| 1. Intensity Tolerance<br>2. *m/z* Tolerance (sample-to-sample)<br>3. Retention Time Tolerance<br>4. Minimum scans | 20.0%<br>0.003 *m/z* or 5.0 ppm<br>0.10 minutes<br>5 |
| **I. Duplicate Peak Filter** | |
| 1. Filter Mode<br>2. *m/z* Tolerance | New Average<br>0.0015 *m/z* or 0.0 ppm |

| | |
|---|---|
| 3. RT Tolerance | 0.05 minutes |
| **J. Feature Filter** | |
| 1. Height<br>2. No. of Data Points | 1.0E5 to 5.0E10<br>5 to 500 |
| **K. Correlation Grouping (metaCorrelate)** | |
| 1. RT Tolerance<br>2. Minimum Feature Height<br>3. Intensity Threshold for Correlation<br>4. Min. Samples Filter<br>a. Min Samples in All<br>b. Min Samples in Group<br>c. Min %-Intensity Overlap<br>d. Exclude Gap-Filled Features<br>5. Feature Shape Correlation<br>a. Min Data Points<br>b. Min Data Points on Edge<br>c. Measure<br>d. Min Feature Shape Correlation | 0.10 min<br>0.0E0<br>0.00<br><br>Max of 1 sample or 0.0%<br>Max of 0 sample or 0.0%<br>60.0%<br>Checked<br><br>5<br>2<br>Pearson<br>85% |
| **L. Ion Identity Networking** | |
| 1. *m/z* Tolerance (intra-sample)<br>2. Check<br>3. Minimum Height<br>4. Ion Identity Library<br>a. Maximum Charge<br>b. Maximum Molecules/Cluster<br>c. Selected Adducts (positive)<br>d. Selected Adducts (negative)<br>5. Annotation Refinement<br>a. Delete Smaller Networks: Link Threshold | 0.0015 *m/z* or 0.0 ppm<br>One Feature<br>0.0E0<br><br>2<br>2<br>$[M+H]^+$, $[M+NH_4]^+$, $[M+Na]^+$, $[M+K]^+$<br>$[M-H]^-$, $[M+Cl]^-$, $[M+FA]^-$, $[M+Acetate]^-$, $[M+Br]^-$<br><br>4 |

**Table S3.** Differential metabolite features identified in MPL-treated HAECs by untargeted LC–MS metabolomics (FDR < 0.05).

| Feature ID | FDR | Features (m/z-Retention time) | Formula | Name | SMILES |
|---|---|---|---|---|---|
| **5899p** | 0.036945 | 528.308-6.34 | C27H46NO7P | LysoPE(22:5(4Z,7Z,10Z,13Z,16Z)/0:0) | CCCCCC=CCC=CCC=CCC=CCC=CCCC(=O)OCC(COP(=O)(O)OCCN)O |
| **8005p** | 0.036959 | 526.314-7.53 | C24H48NO9P | 1-Stearoylglycerophosphoserine | CCCCCCCCCCCCCCCCCC(=O)OCC(COP(=O)(O)OCC(C(=O)O)N)O |
| **5258p** | 0.038136 | 223.112-5.67 | C16H14O | (3E)-3,4-Diphenylbut-3-en-2-one | CC(=O)C(=CC1=CC=CC=C1)C2=CC=CC=C2 |
| **4406p** | 5.89E-05 | 241.122-4.41 | C16H16O2 | (E)-alpha,beta-Dimethoxystilbene | COC(=C(C1=CC=CC=C1)OC)C2=CC=CC=C2 |
| **6882p** | 0.009713 | 508.304-6.97 | C24H46NO8P | [(2R)-1-[2-aminoethoxy(hydroxy)phosphoryl]oxy-3-heptanoyloxypropan-2-yl] (E)-dodec-9-enoate | CCCCCCC(=O)OCC(COP(=O)(O)OCCN)OC(=O)CCCCCCCC=CCC |
| **5675p** | 0.000304 | 480.272-6.16 | C22H42NO8P | [(2R)-1-[2-aminoethoxy(hydroxy)phosphoryl]oxy-3-heptanoyloxypropan-2-yl] dec-9-enoate | CCCCCCC(=O)OCC(COP(=O)(O)OCCN)OC(=O)CCCCCCCC=C |
| **5796p** | 0.009805 | 480.272-6.25 | C22H42NO8P | [(2R)-1-[2-aminoethoxy(hydroxy)phosphoryl]oxy-3-heptanoyloxypropan-2-yl] dec-9-enoate | CCCCCCC(=O)OCC(COP(=O)(O)OCCN)OC(=O)CCCCCCCC=C |
| **5812p** | 0.016336 | 522.319-6.27 | C25H48NO8P | [(2R)-1-[2-aminoethoxy(hydroxy)phosphoryl]oxy-3-nonanoyloxypropan-2-yl] (Z)-undec-9-enoate | CCCCCCCCC(=O)OCC(COP(=O)(O)OCCN)OC(=O)CCCCCCCC=CC |

| | | | | | |
|---|---|---|---|---|---|
| **8033p** | 0.033794 | 560.335-7.54 | C28H50NO8P | [1-[2-aminoethoxy(hydroxy)phosphoryl]oxy-3-heptanoyloxypropan-2-yl] (7Z,10Z,13Z)-hexadeca-7,10,13-trienoate | CCCCCCC(=O)OCC(COP(=O)(O)OCCN)OC(=O)CCCCCC=CCC=CCC=CCC |
| **6014p** | 0.030299 | 572.335-6.42 | C29H50NO8P | [2-[(4Z,7Z,10Z,13Z)-hexadeca-4,7,10,13-tetraenoyl]oxy-3-pentanoyloxypropyl] 2-(trimethylazaniumyl)ethyl phosphate | CCCCC(=O)OCC(COP(=O)(O)OCC[N+](C)(C)C)OC(=O)CCC=CCC=CCC=CCC=CCC |
| **8256p** | 0.02433 | 650.44-7.63 | C33H64NO9P | [3-Hexadecanoyloxy-2-(9-oxononanoyloxy)propyl] 2-(trimethylazaniumyl)ethyl phosphate | CCCCCCCCCCCCCCCC(=O)OCC(COP(=O)(O)OCC[N+](C)(C)C)OC(=O)CCCCCCCC=O |
| **7660p** | 0.033794 | 636.424-7.37 | C32H62NO9P | [O-[1-O-Palmitoyl-2-O-(8-oxooctanoyl)-L-glycero-3-phospho]choline]anion | CCCCCCCCCCCCCCCC(=O)OCC(CO[P+](O)([O-])OCC[N+](C)(C)C)OC(=O)CCCCCCC=O |
| **5600p** | 0.022357 | 365.115-6.07 | C21H12N6O | 1-(3-Cyano-4-phenylpyrazole-1-carbonyl)-4-phenylpyrazole-3-carbonitrile | C1=CC=C(C=C1)C2=CN(N=C2C#N)C(=O)N3C=C(C(=N3)C#N)C4=CC=CC=C4 |
| **6085p** | 0.033867 | 417.336-6.46 | C27H44O3 | 1-(5-hydroperoxy-6-methylhept-6-en-2-yl)-9a,11a-dimethyl-1H,2H,3H,3aH,3bH,4H,6H,7H,8H,9H,9aH,9bH,10H,11H,11aH-cyclopenta[a]phenanthren-7-ol | CC(CCC(C(=C)C)OO)C1CCC2C1(CCC3C2CC=C4C3(CCC(C4)O)C)C |
| **2100p** | 0.022147 | 264.159-2.29 | C15H21NO3 | 1-[4-(2-Morpholin-4-ylethoxy)phenyl]propan-1-one | CCC(=O)C1=CC=C(C=C1)OCCN2CCOCC2 |
| **5689p** | 0.02771 | 502.293-6.17 | C25H44NO7P | PE(20:4(5Z,8Z,11Z,14Z)/0:0) | CCCCCC=CCC=CCC=CCC=CCCCC(=O)OCC(COP(=O)(O)OCCN)O |
| **6185p** | 0.034519 | 454.293-6.51 | C21H44NO7P | 1-Hexadecanoyl-sn-glycero-3-phosphoethanolamine | CCCCCCCCCCCCCCCC(=O)OCC(COP(=O)(O)OCCN)O |
| **5488p** | 0.01031 | 496.303-5.96 | C23H46NO8P | 1-Octanoyl-2-heptanoylphosphatidylcholine | CCCCCCCC(=O)OCC(COP(=O)(O)OCC[N+](C)(C)C)OC(=O)CCCCCC |
| **6491p** | 0.043405 | 480.309-6.71 | C23H46NO7P | 1-Oleoyl-Sn-Glycero-3-Phosphoethanolamine | CCCCCCCCC=CCCCCCCCC(=O)OCC(COP(=O)(O)OCCN)O |
| **6632p** | 0.007097 | 524.334-6.8 | C25H50NO8P | 1-Pentadecanoyl-2-acetyl-sn-glycero-3-phosphocholine | CCCCCCCCCCCCCCC(=O)OCC(COP(=O)(O)OCC[N+](C)(C)C)OC(=O)C |
| **6193p** | 0.008028 | 225.091-6.52 | C15H12O2 | 1,3-diphenylpropane-1,3-dione | C1=CC=C(C=C1)C(=O)CC(=O)C2=CC=CC=C2 |
| **5896p** | 0.0072 | 329.154-6.34 | C23H20O2 | 1,3,5-Triphenyl-1,5-pentanedione | C1=CC=C(C=C1)C(CC(=O)C2=CC=CC=C2)CC(=O)C3=CC=CC=C3 |
| **6910p** | 0.033887 | 311.18-6.99 | C24H22 | 1,4-Bis(2-phenylprop-1-enyl)benzene | CC(=CC1=CC=C(C=C1)C=C(C)C2=CC=CC=C2)C3=CC=CC=C3 |
| **6488p** | 0.014803 | 532.303-6.71 | C28H41N3O7 | 18-Amino-4,5-dehydro-17-demethylreblastatin | CC1CC(C(C(C=C(C(C(C=CC=C(C(=O)NC2=CC(=C(C(=C2)N)O)C1)C)OC)OC(=O)N)C)C)O)OC |
| **6793p** | 0.023031 | 532.304-6.91 | C28H41N3O7 | 18-Amino-4,5-dehydro-17-demethylreblastatin | CC1CC(C(C(C=C(C(C(C=CC=C(C(=O)NC2=CC(=C(C(=C2)N)O)C1)C)OC)OC(=O)N)C)C)O)OC |
| **3780p** | 0.010811 | 260.201-3.81 | C17H25NO | 1beta-Phenyl-3alpha-piperidinocyclohexanol | C1CCN(CC1)C2CCCC(C2)(C3=CC=CC=C3)O |
| **6905p** | 0.005789 | 608.392-6.99 | C30H58NO9P | 2-[[3-Heptadecanoyloxy-2-(5-oxopentanoyloxy)propoxy]-hydroxyphosphoryl]oxyethyl-trimethylazanium | CCCCCCCCCCCCCCCCC(=O)OCC(COP(=O)(O)OCC[N+](C)(C)C)OC(=O)CCCC=O |

| | | | | | |
|---|---|---|---|---|---|
| **7221p** | 0.014803 | 608.393-7.15 | C30H58NO9P | 2-[[3-Heptadecanoyloxy-2-(5-oxopentanoyloxy)propoxy]-hydroxyphosphoryl]oxyethyl-trimethylazanium | CCCCCCCCCCCCCCCCC(=O)OCC(COP(=O)(O)OCC[N+](C)(C)C)OC(=O)CCCC=O |
| **6001p** | 0.021933 | 530.288-6.41 | C29H35N7O3 | 2-[2-[2-methoxy-4-(4-methylpiperazin-1-yl)anilino]pyrimidin-4-yl]-2-methyl-N-[3-(prop-2-enoylamino)phenyl]propanamide | CC(C)(C1=NC(=NC=C1)NC2=C(C=C(C=C2)N3CCN(CC3)C)OC)C(=O)NC4=CC=CC(=C4)NC(=O)C=C |
| **6897p** | 0.018127 | 550.35-6.98 | C27H52NO8P | 2-[hydroxy-[(2S)-3-octanoyloxy-2-[(E)-undec-9-enoyl]oxypropoxy]phosphoryl]oxyethyl-trimethylazanium | CCCCCCCC(=O)OCC(COP(=O)(O)OCC[N+](C)(C)C)OC(=O)CCCCCCCC=CC |
| **5037p** | 0.007429 | 239.107-5.29 | C16H14O2 | 2,3-Diphenyl-2-butenoic acid | CC(=C(C1=CC=CC=C1)C(=O)O)C2=CC=CC=C2 |
| **794p** | 0.033887 | 330.073-0.59 | C15H11N3O6 | 3-[[2-(4-Nitroanilino)-2-oxoacetyl]amino]benzoic acid | C1=CC(=CC(=C1)NC(=O)C(=O)NC2=CC=C(C=C2)[N+](=O)[O-])C(=O)O |
| **1420p** | 0.01437 | 127.039-1.16 | C6H6O3 | 3-Methyl-4-hydroxyfuran-2-carbaldehyde | CC1=C(OC=C1O)C=O |
| **2375p** | 7.63E-05 | 174.185-2.67 | C10H23NO | 4-(Hexylamino)butan-1-OL | CCCCCCNCCCCO |
| **1551p** | 0.001157 | 169.032-1.49 | C8H8O2S | 4-[(R)-methylsulfinyl]benzaldehyde | CS(=O)C1=CC=C(C=C1)C=O |
| **6277p** | 0.023378 | 556.304-6.56 | C30H41N3O7 | Cbz-Ala-Gly-Tyr(tBu)-OtBu | CC(C(=O)NCC(=O)NC(CC1=CC=C(C=C1)OC(C)(C)C)C(=O)OC(C)(C)C)NC(=O)OCC2=CC=CC=C2 |
| **5339p** | 0.00752 | 209.096-5.77 | C15H12O | Chalcone | C1=CC=C(C=C1)C=CC(=O)C2=CC=CC=C2 |
| **704p** | 0.022126 | 184.073-0.57 | C5H14NO4P | phosphocholine | C[N+](C)(C)CCOP(=O)(O)O |
| **674p** | 0.021365 | 142.948-0.56 | C4H3AsO | Furylarsine | C1=COC(=C1)[As] |
| **6589p** | 0.003062 | 482.288-6.78 | C25H35N7O3 | N-(2-morpholin-4-ylethyl)-6-[4-[3-(propan-2-ylamino)pyridin-2-yl]piperazine-1-carbonyl]pyridine-3-carboxamide | CC(C)NC1=C(N=CC=C1)N2CCN(CC2)C(=O)C3=NC=C(C=C3)C(=O)NCCN4CCOCC4 |
| **6954p** | 0.022126 | 558.32-7.01 | C30H43N3O7 | N-[(2R)-1-[[(2S)-1-[[(2S)-3-cyclobutyl-1-[(2R)-2-methyloxiran-2-yl]-1-oxopropan-2-yl]amino]-3-(4-methoxyphenyl)-1-oxopropan-2-yl]amino]-1-oxopropan-2-yl]-4-hydroxycyclohexane-1-carboxamide | CC(C(=O)NC(CC1=CC=C(C=C1)OC)C(=O)NC(CC2CCC2)C(=O)C3(CO3)C)NC(=O)C4CCC(CC4)O |
| **507p** | 0.043012 | 206.055-0.53 | C6H11ClF3NO | N-[2-(2-chloroethoxy)ethyl]-2,2,2-trifluoroethanamine | C(COCCCl)NCC(F)(F)F |
| **5928p** | 0.036636 | 554.288-6.35 | C31H35N7O3 | N-[4-(2-amino-1H-imidazol-5-yl)cyclohexyl]-2-(2,2-diphenylethyl)-6-methyl-1,3-dioxo-5,8-dihydro-[1,2,4]triazolo[1,2-a]pyridazine-8-carboxamide | CC1=CC(N2C(=O)N(C(=O)N2C1)CC(C3=CC=CC=C3)C4=CC=CC=C4)C(=O)NC5CCC(CC5)C6=CN=C(N6)N |
| **2808p** | 0.029138 | 225.139-3.02 | C15H16N2 | n-Butyl-b-carboline | CCCCC1=NC=CC2=C1NC3=CC=CC=C23 |
| **5885p** | 0.007097 | 351.135-6.33 | C25H18O2 | Phenyl-[4-(4-phenylphenoxy)phenyl]methanone | C1=CC=C(C=C1)C2=CC=C(C=C2)OC3=CC=C(C=C3)C(=O)C4=CC=CC=C4 |
| **4407p** | 3.63E-05 | 223.112-4.41 | C16H14O | Phenyl[4-(prop-1-en-2-yl)phenyl]methanone | CC(=C)C1=CC=C(C=C1)C(=O)C2=CC=CC=C2 |
| **4569p** | 0.002676 | 223.111-4.66 | C16H14O | Phenyl[4-(prop-1-en-2-yl)phenyl]methanone | CC(=C)C1=CC=C(C=C1)C(=O)C2=CC=CC=C2 |

**Table S2.** Parameters used of Mzmine peak picking analysis of MS raw data

| Module | Parameters Used |
|---|---|
| **A. Mass Detection (first, for MS1 data)** | |
| 1. MS Level Filter<br>2. Polarity<br><br>3. Spectrum Type<br>4. Mass Detector<br>5. Noise Level | MS1, level 1<br>positive and negative<br>(processed in separate batches)<br>centroid<br>centroid<br>5.0E3 |
| **B. Mass Detection (second, for MS2 data)** | |
| 1. MS Level Filter<br>2. Polarity<br><br>3. Spectrum Type<br>4. Mass Detector<br>5. Noise Level | MS2, level 2<br>positive and negative<br>(processed in separate batches)<br>centroid<br>centroid<br>100 |
| **C. Chromatogram Builder** | |
| 1. MS Level Filter<br>2. Polarity<br><br>3. Spectrum Type<br>4. Min. Consecutive Scans<br>5. Min. Intensity for Consecutive Scans<br>6. Min. Absolute Height<br>7. *m/z* Tolerance (scan-to-scan) | MS1, level 1<br>positive and negative<br>(processed in separate batches)<br>centroid<br>5<br>5.0E3<br>1.0E5<br>0.003 *m/z* or 5.0 ppm |
| **D. Local Minimum Feature Resolver** | |
| 1. MS1 to MS2 Precursor Tolerance (*m/z*)<br>2. Retention Time Filter<br>3. Minimum Relative Feature Height<br>4. Dimension<br>5. Chromatographic Threshold<br>6. Minimum Search Range RT<br>7. Minimum Absolute Height<br>8. Min. Ratio of Peak Top/Edge<br>9. Peak Duration Range<br>10. Minimum Scans (data points) | 0.003 m/z or 5.0 ppm<br>Use Tolerance, 0.20 min.<br>25%<br>Retention Time<br>70%<br>0.05<br>1.0E5<br>1.50<br>0.00 to 5.00<br>5 |
| **E. $^{13}C$ Isotope Filter** | |
| 1. *m/z* Tolerance (intra-sample)<br>2. Retention Time Tolerance<br>3. Monotonic Shape<br>4. Maximum Charge<br>5. Representative Isotope<br>6. Never Remove Feature with MS2 | 0.0015 *m/z* or 0.0 ppm<br>0.05 min<br>Checked<br>2<br>Most Intense<br>Checked |
| **F. Isotopic Peaks Finder** | |
| 1. Chemical Elements<br>2. *m/z* Tolerance (feature-to-scan)<br>3. Maximum Charge of Isotope *m/z*<br>4. Search in Scans | H, C, N, O, S, Cl, Br, P, F<br>0.002 *m/z* or 0.0 ppm<br>2<br>Single Most Intense |
| **G. Join Aligner** | |
| 1. *m/z* Tolerance (sample-to-sample)<br>2. Weight for *m/z*<br>3. Retention Time Tolerance<br>4. Weight for RT<br>5. Mobility Weight | 0.003 *m/z* or 5.0 ppm<br>2<br>0.10 minutes<br>1<br>1.0 |
| **H. Peak Finder (Gap Filling)** | |
| 1. Intensity Tolerance<br>2. *m/z* Tolerance (sample-to-sample)<br>3. Retention Time Tolerance<br>4. Minimum scans | 20.0%<br>0.003 *m/z* or 5.0 ppm<br>0.10 minutes<br>5 |
| **I. Duplicate Peak Filter** | |
| 1. Filter Mode<br>2. *m/z* Tolerance | New Average<br>0.0015 *m/z* or 0.0 ppm |

| | |
|---|---|
| 3. RT Tolerance | 0.05 minutes |
| **J. Feature Filter** | |
| 1. Height<br>2. No. of Data Points | 1.0E5 to 5.0E10<br>5 to 500 |
| **K. Correlation Grouping (metaCorrelate)** | |
| 1. RT Tolerance<br>2. Minimum Feature Height<br>3. Intensity Threshold for Correlation<br>4. Min. Samples Filter<br>a. Min Samples in All<br>b. Min Samples in Group<br>c. Min %-Intensity Overlap<br>d. Exclude Gap-Filled Features<br>5. Feature Shape Correlation<br>a. Min Data Points<br>b. Min Data Points on Edge<br>c. Measure<br>d. Min Feature Shape Correlation | 0.10 min<br>0.0E0<br>0.00<br><br>Max of 1 sample or 0.0%<br>Max of 0 sample or 0.0%<br>60.0%<br>Checked<br><br>5<br>2<br>Pearson<br>85% |
| **L. Ion Identity Networking** | |
| 1. *m/z* Tolerance (intra-sample)<br>2. Check<br>3. Minimum Height<br>4. Ion Identity Library<br>a. Maximum Charge<br>b. Maximum Molecules/Cluster<br>c. Selected Adducts (positive)<br>d. Selected Adducts (negative)<br>5. Annotation Refinement<br>a. Delete Smaller Networks: Link Threshold | 0.0015 *m/z* or 0.0 ppm<br>One Feature<br>0.0E0<br><br>2<br>2<br>$[M+H]^+$, $[M+NH_4]^+$, $[M+Na]^+$, $[M+K]^+$<br>$[M-H]^-$, $[M+Cl]^-$, $[M+FA]^-$, $[M+Acetate]^-$, $[M+Br]^-$<br><br>4 |

**Table S3.** Differential metabolite features identified in MPL-treated HAECs by untargeted LC–MS metabolomics (FDR < 0.05).

| Feature ID | FDR | Features (m/z-Retention time) | Formula | Name | SMILES |
|---|---|---|---|---|---|
| **5899p** | 0.036945 | 528.308-6.34 | C27H46NO7P | LysoPE(22:5(4Z,7Z,10Z,13Z,16Z)/0:0) | CCCCCC=CCC=CCC=CCC=CCC=CCCC(=O)OCC(COP(=O)(O)OCCN)O |
| **8005p** | 0.036959 | 526.314-7.53 | C24H48NO9P | 1-Stearoylglycerophosphoserine | CCCCCCCCCCCCCCCCCC(=O)OCC(COP(=O)(O)OCC(C(=O)O)N)O |
| **5258p** | 0.038136 | 223.112-5.67 | C16H14O | (3E)-3,4-Diphenylbut-3-en-2-one | CC(=O)C(=CC1=CC=CC=C1)C2=CC=CC=C2 |
| **4406p** | 5.89E-05 | 241.122-4.41 | C16H16O2 | (E)-alpha,beta-Dimethoxystilbene | COC(=C(C1=CC=CC=C1)OC)C2=CC=CC=C2 |
| **6882p** | 0.009713 | 508.304-6.97 | C24H46NO8P | [(2R)-1-[2-aminoethoxy(hydroxy)phosphoryl]oxy-3-heptanoyloxypropan-2-yl] (E)-dodec-9-enoate | CCCCCCC(=O)OCC(COP(=O)(O)OCCN)OC(=O)CCCCCCCC=CCC |
| **5675p** | 0.000304 | 480.272-6.16 | C22H42NO8P | [(2R)-1-[2-aminoethoxy(hydroxy)phosphoryl]oxy-3-heptanoyloxypropan-2-yl] dec-9-enoate | CCCCCCC(=O)OCC(COP(=O)(O)OCCN)OC(=O)CCCCCCCC=C |
| **5796p** | 0.009805 | 480.272-6.25 | C22H42NO8P | [(2R)-1-[2-aminoethoxy(hydroxy)phosphoryl]oxy-3-heptanoyloxypropan-2-yl] dec-9-enoate | CCCCCCC(=O)OCC(COP(=O)(O)OCCN)OC(=O)CCCCCCCC=C |
| **5812p** | 0.016336 | 522.319-6.27 | C25H48NO8P | [(2R)-1-[2-aminoethoxy(hydroxy)phosphoryl]oxy-3-nonanoyloxypropan-2-yl] (Z)-undec-9-enoate | CCCCCCCCC(=O)OCC(COP(=O)(O)OCCN)OC(=O)CCCCCCCC=CC |

| | | | | | |
|---|---|---|---|---|---|
| **8033p** | 0.033794 | 560.335-7.54 | C28H50NO8P | [1-[2-aminoethoxy(hydroxy)phosphoryl]oxy-3-heptanoyloxypropan-2-yl] (7Z,10Z,13Z)-hexadeca-7,10,13-trienoate | CCCCCCC(=O)OCC(COP(=O)(O)OCCN)OC(=O)CCCCCC=CCC=CCC=CCC |
| **6014p** | 0.030299 | 572.335-6.42 | C29H50NO8P | [2-[(4Z,7Z,10Z,13Z)-hexadeca-4,7,10,13-tetraenoyl]oxy-3-pentanoyloxypropyl] 2-(trimethylazaniumyl)ethyl phosphate | CCCCC(=O)OCC(COP(=O)(O)OCC[N+](C)(C)C)OC(=O)CCC=CCC=CCC=CCC=CCC |
| **8256p** | 0.02433 | 650.44-7.63 | C33H64NO9P | [3-Hexadecanoyloxy-2-(9-oxononanoyloxy)propyl] 2-(trimethylazaniumyl)ethyl phosphate | CCCCCCCCCCCCCCCC(=O)OCC(COP(=O)(O)OCC[N+](C)(C)C)OC(=O)CCCCCCCC=O |
| **7660p** | 0.033794 | 636.424-7.37 | C32H62NO9P | [O-[1-O-Palmitoyl-2-O-(8-oxooctanoyl)-L-glycero-3-phospho]choline]anion | CCCCCCCCCCCCCCCC(=O)OCC(CO[P+](O)([O-])OCC[N+](C)(C)C)OC(=O)CCCCCCC=O |
| **5600p** | 0.022357 | 365.115-6.07 | C21H12N6O | 1-(3-Cyano-4-phenylpyrazole-1-carbonyl)-4-phenylpyrazole-3-carbonitrile | C1=CC=C(C=C1)C2=CN(N=C2C#N)C(=O)N3C=C(C(=N3)C#N)C4=CC=CC=C4 |
| **6085p** | 0.033867 | 417.336-6.46 | C27H44O3 | 1-(5-hydroperoxy-6-methylhept-6-en-2-yl)-9a,11a-dimethyl-1H,2H,3H,3aH,3bH,4H,6H,7H,8H,9H,9aH,9bH,10H,11H,11aH-cyclopenta[a]phenanthren-7-ol | CC(CCC(C(=C)C)OO)C1CCC2C1(CCC3C2CC=C4C3(CCC(C4)O)C)C |
| **2100p** | 0.022147 | 264.159-2.29 | C15H21NO3 | 1-[4-(2-Morpholin-4-ylethoxy)phenyl]propan-1-one | CCC(=O)C1=CC=C(C=C1)OCCN2CCOCC2 |
| **5689p** | 0.02771 | 502.293-6.17 | C25H44NO7P | PE(20:4(5Z,8Z,11Z,14Z)/0:0) | CCCCCC=CCC=CCC=CCC=CCCCC(=O)OCC(COP(=O)(O)OCCN)O |
| **6185p** | 0.034519 | 454.293-6.51 | C21H44NO7P | 1-Hexadecanoyl-sn-glycero-3-phosphoethanolamine | CCCCCCCCCCCCCCCC(=O)OCC(COP(=O)(O)OCCN)O |
| **5488p** | 0.01031 | 496.303-5.96 | C23H46NO8P | 1-Octanoyl-2-heptanoylphosphatidylcholine | CCCCCCCC(=O)OCC(COP(=O)(O)OCC[N+](C)(C)C)OC(=O)CCCCCC |
| **6491p** | 0.043405 | 480.309-6.71 | C23H46NO7P | 1-Oleoyl-Sn-Glycero-3-Phosphoethanolamine | CCCCCCCCC=CCCCCCCCC(=O)OCC(COP(=O)(O)OCCN)O |
| **6632p** | 0.007097 | 524.334-6.8 | C25H50NO8P | 1-Pentadecanoyl-2-acetyl-sn-glycero-3-phosphocholine | CCCCCCCCCCCCCCC(=O)OCC(COP(=O)(O)OCC[N+](C)(C)C)OC(=O)C |
| **6193p** | 0.008028 | 225.091-6.52 | C15H12O2 | 1,3-diphenylpropane-1,3-dione | C1=CC=C(C=C1)C(=O)CC(=O)C2=CC=CC=C2 |
| **5896p** | 0.0072 | 329.154-6.34 | C23H20O2 | 1,3,5-Triphenyl-1,5-pentanedione | C1=CC=C(C=C1)C(CC(=O)C2=CC=CC=C2)CC(=O)C3=CC=CC=C3 |
| **6910p** | 0.033887 | 311.18-6.99 | C24H22 | 1,4-Bis(2-phenylprop-1-enyl)benzene | CC(=CC1=CC=C(C=C1)C=C(C)C2=CC=CC=C2)C3=CC=CC=C3 |
| **6488p** | 0.014803 | 532.303-6.71 | C28H41N3O7 | 18-Amino-4,5-dehydro-17-demethylreblastatin | CC1CC(C(C(C=C(C(C(C=CC=C(C(=O)NC2=CC(=C(C(=C2)N)O)C1)C)OC)OC(=O)N)C)C)O)OC |
| **6793p** | 0.023031 | 532.304-6.91 | C28H41N3O7 | 18-Amino-4,5-dehydro-17-demethylreblastatin | CC1CC(C(C(C=C(C(C(C=CC=C(C(=O)NC2=CC(=C(C(=C2)N)O)C1)C)OC)OC(=O)N)C)C)O)OC |
| **3780p** | 0.010811 | 260.201-3.81 | C17H25NO | 1beta-Phenyl-3alpha-piperidinocyclohexanol | C1CCN(CC1)C2CCCC(C2)(C3=CC=CC=C3)O |
| **6905p** | 0.005789 | 608.392-6.99 | C30H58NO9P | 2-[[3-Heptadecanoyloxy-2-(5-oxopentanoyloxy)propoxy]-hydroxyphosphoryl]oxyethyl-trimethylazanium | CCCCCCCCCCCCCCCCC(=O)OCC(COP(=O)(O)OCC[N+](C)(C)C)OC(=O)CCCC=O |

| | | | | | |
|---|---|---|---|---|---|
| **7221p** | 0.014803 | 608.393-7.15 | C30H58NO9P | 2-[[3-Heptadecanoyloxy-2-(5-oxopentanoyloxy)propoxy]-hydroxyphosphoryl]oxyethyl-trimethylazanium | CCCCCCCCCCCCCCCCC(=O)OCC(COP(=O)(O)OCC[N+](C)(C)C)OC(=O)CCCC=O |
| **6001p** | 0.021933 | 530.288-6.41 | C29H35N7O3 | 2-[2-[2-methoxy-4-(4-methylpiperazin-1-yl)anilino]pyrimidin-4-yl]-2-methyl-N-[3-(prop-2-enoylamino)phenyl]propanamide | CC(C)(C1=NC(=NC=C1)NC2=C(C=C(C=C2)N3CCN(CC3)C)OC)C(=O)NC4=CC=CC(=C4)NC(=O)C=C |
| **6897p** | 0.018127 | 550.35-6.98 | C27H52NO8P | 2-[hydroxy-[(2S)-3-octanoyloxy-2-[(E)-undec-9-enoyl]oxypropoxy]phosphoryl]oxyethyl-trimethylazanium | CCCCCCCC(=O)OCC(COP(=O)(O)OCC[N+](C)(C)C)OC(=O)CCCCCCCC=CC |
| **5037p** | 0.007429 | 239.107-5.29 | C16H14O2 | 2,3-Diphenyl-2-butenoic acid | CC(=C(C1=CC=CC=C1)C(=O)O)C2=CC=CC=C2 |
| **794p** | 0.033887 | 330.073-0.59 | C15H11N3O6 | 3-[[2-(4-Nitroanilino)-2-oxoacetyl]amino]benzoic acid | C1=CC(=CC(=C1)NC(=O)C(=O)NC2=CC=C(C=C2)[N+](=O)[O-])C(=O)O |
| **1420p** | 0.01437 | 127.039-1.16 | C6H6O3 | 3-Methyl-4-hydroxyfuran-2-carbaldehyde | CC1=C(OC=C1O)C=O |
| **2375p** | 7.63E-05 | 174.185-2.67 | C10H23NO | 4-(Hexylamino)butan-1-OL | CCCCCCNCCCCO |
| **1551p** | 0.001157 | 169.032-1.49 | C8H8O2S | 4-[(R)-methylsulfinyl]benzaldehyde | CS(=O)C1=CC=C(C=C1)C=O |
| **6277p** | 0.023378 | 556.304-6.56 | C30H41N3O7 | Cbz-Ala-Gly-Tyr(tBu)-OtBu | CC(C(=O)NCC(=O)NC(CC1=CC=C(C=C1)OC(C)(C)C)C(=O)OC(C)(C)C)NC(=O)OCC2=CC=CC=C2 |
| **5339p** | 0.00752 | 209.096-5.77 | C15H12O | Chalcone | C1=CC=C(C=C1)C=CC(=O)C2=CC=CC=C2 |
| **704p** | 0.022126 | 184.073-0.57 | C5H14NO4P | phosphocholine | C[N+](C)(C)CCOP(=O)(O)O |
| **674p** | 0.021365 | 142.948-0.56 | C4H3AsO | Furylarsine | C1=COC(=C1)[As] |
| **6589p** | 0.003062 | 482.288-6.78 | C25H35N7O3 | N-(2-morpholin-4-ylethyl)-6-[4-[3-(propan-2-ylamino)pyridin-2-yl]piperazine-1-carbonyl]pyridine-3-carboxamide | CC(C)NC1=C(N=CC=C1)N2CCN(CC2)C(=O)C3=NC=C(C=C3)C(=O)NCCN4CCOCC4 |
| **6954p** | 0.022126 | 558.32-7.01 | C30H43N3O7 | N-[(2R)-1-[[(2S)-1-[[(2S)-3-cyclobutyl-1-[(2R)-2-methyloxiran-2-yl]-1-oxopropan-2-yl]amino]-3-(4-methoxyphenyl)-1-oxopropan-2-yl]amino]-1-oxopropan-2-yl]-4-hydroxycyclohexane-1-carboxamide | CC(C(=O)NC(CC1=CC=C(C=C1)OC)C(=O)NC(CC2CCC2)C(=O)C3(CO3)C)NC(=O)C4CCC(CC4)O |
| **507p** | 0.043012 | 206.055-0.53 | C6H11ClF3NO | N-[2-(2-chloroethoxy)ethyl]-2,2,2-trifluoroethanamine | C(COCCCl)NCC(F)(F)F |
| **5928p** | 0.036636 | 554.288-6.35 | C31H35N7O3 | N-[4-(2-amino-1H-imidazol-5-yl)cyclohexyl]-2-(2,2-diphenylethyl)-6-methyl-1,3-dioxo-5,8-dihydro-[1,2,4]triazolo[1,2-a]pyridazine-8-carboxamide | CC1=CC(N2C(=O)N(C(=O)N2C1)CC(C3=CC=CC=C3)C4=CC=CC=C4)C(=O)NC5CCC(CC5)C6=CN=C(N6)N |
| **2808p** | 0.029138 | 225.139-3.02 | C15H16N2 | n-Butyl-b-carboline | CCCCC1=NC=CC2=C1NC3=CC=CC=C23 |
| **5885p** | 0.007097 | 351.135-6.33 | C25H18O2 | Phenyl-[4-(4-phenylphenoxy)phenyl]methanone | C1=CC=C(C=C1)C2=CC=C(C=C2)OC3=CC=C(C=C3)C(=O)C4=CC=CC=C4 |
| **4407p** | 3.63E-05 | 223.112-4.41 | C16H14O | Phenyl[4-(prop-1-en-2-yl)phenyl]methanone | CC(=C)C1=CC=C(C=C1)C(=O)C2=CC=CC=C2 |
| **4569p** | 0.002676 | 223.111-4.66 | C16H14O | Phenyl[4-(prop-1-en-2-yl)phenyl]methanone | CC(=C)C1=CC=C(C=C1)C(=O)C2=CC=CC=C2 |
